# Anomalous reverse transcription through chemical modifications in polyadenosine stretches

**DOI:** 10.1101/2020.01.07.897843

**Authors:** Wipapat Kladwang, Ved V. Topkar, Bei Liu, Tracy L. Hodges, Sarah C. Keane, Hashim al-Hashimi, Rhiju Das

**Affiliations:** Department of Biochemistry, Stanford University School of Medicine, Stanford CA 94305; Biophysics Program, Stanford University, Stanford CA 94305; Department of Biochemistry, Duke University School of Medicine, Durham NC 27710; Biophysics Program, University of Michigan, Ann Arbor MI 48109; Department of Chemistry, University of Michigan, Ann Arbor MI 48109; Department of Chemistry, Duke University School of Medicine, Durham NC 27710; Department of Physics, Stanford University, Stanford CA 94305

## Abstract

Thermostable reverse transcriptases are workhorse enzymes underlying nearly all modern techniques for RNA structure mapping and for transcriptome-wide discovery of RNA chemical modifications. Despite their wide use, these enzymes’ behaviors at chemical modified nucleotides remain poorly understood. Wellington-Oguri et al. recently reported an apparent loss of chemical modification within putatively unstructured polyadenosine stretches modified by dimethyl sulfate or 2’ hydroxyl acylation, as probed by reverse transcription. Here, re-analysis of these and other publicly available data, capillary electrophoresis experiments on chemically modified RNAs, and nuclear magnetic resonance spectroscopy on A_12_ and variants show that this effect is unlikely to arise from an unusual structure of polyadenosine. Instead, tests of different reverse transcriptases on chemically modified RNAs and molecules synthesized with single 1-methyladenosines implicate a previously uncharacterized reverse transcriptase behavior: near-quantitative bypass through chemical modifications within polyadenosine stretches. All tested natural and engineered reverse transcriptases (MMLV; SuperScript II, III, and IV; TGIRT-III; and MarathonRT) exhibit this anomalous bypass behavior. Accurate DMS-guided structure modeling of the polyadenylated HIV-1 3’ untranslated region RNA requires taking into account this anomaly. Our results suggest that poly(rA-dT) hybrid duplexes can trigger unexpectedly effective reverse transcriptase bypass and that chemical modifications in poly(A) mRNA tails may be generally undercounted.

## Introduction

RNA molecules play extensive roles in gene regulation and expression in all forms of life.^1^ Recent years have seen an explosion of research into the chemical modification of non-coding RNA, mRNA, and synthetic RNA molecules, although the positions and modification rates of bases like m^1^A (1-methyladenosine) in human cells remain under intense debate.^2-7^ In parallel to these ‘epitranscriptomic’ studies, researchers interested in how RNA molecules fold into elaborate structures measure the reactivity of RNA nucleotides to chemical modification reagents to probe their structural accessibility.^8^ To read out the chemical modifications, most modern technologies take advantage of reverse transcriptase (RTase) enzymes either terminating at modified nucleotides or reading through these positions and leaving mutations, insertions, or deletions in the complementary DNA (cDNA) that correspond to the site of RNA modification. Sequencing of the resulting cDNAs then affords single-nucleotide-resolution measurements of chemical modifications of the original RNA samples.^2, 7, 9-10^

RTase-based mapping has begun to yield large databases of experimental results, such as the RNA Mapping Database (RMDB) of structure mapping profiles, which catalogues over 50,000 sequences.^11^ A surprising finding from these data has been a striking chemical accessibility signature for long stretches of adenosines (A’s).^12^ On one hand, the last (3’-most) six adenosines in such long poly(A) stretches give clear signals for modification by chemical reagents. On the other hand, adenosines 5’ to these last adenosines show striking loss of signal. The observation of this apparent protection pattern includes not only experiments with dimethyl sulfate (DMS) – which methylates chemically accessible N1 positions of adenosines to m^1^A– but also experiments with reagents used for selective 2’-hydroxyl acylation with primer extension (SHAPE), which marks flexible nucleotides (Fig. 1a).^13^ Substitution of either C, G, or U at any A position results in restoration of chemical modification signals for 6 nucleotides upstream (5’) but not downstream (3’) of the substitution (Fig. 1b).

**Fig. 1.**
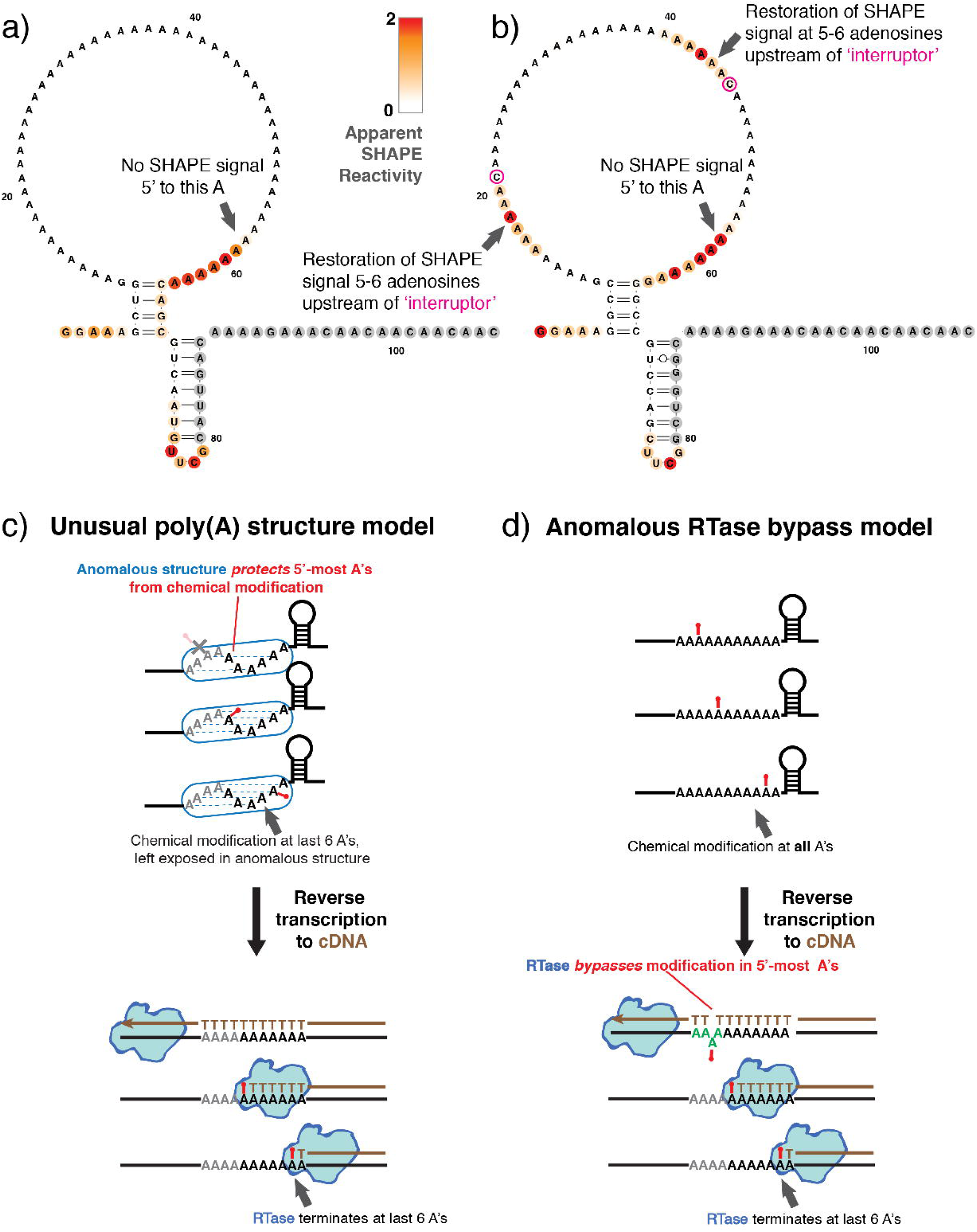
SHAPE mapping profiles for representative poly(A)-containing sequences can be explained by two possible models. (a) SHAPE signal is diminished upstream (5’) of 6 modified adenosines. (b) Installation of cytidine interrupters (magenta) restores SHAPE signal for the 6 adenosines upstream (5’) to the interrupter. Data are from experiments that modified RNA with 1M7 (1-methyl-7-nitroisatoic anhydride) and correspond to RNA Mapping Database entry ETERNA_R69_0000, sequence numbers 851 and 852; data normalized so that GAGUA pentaloops in co-loaded reference constructs give mean SHAPE reactivities of unity.^37^ (c,d) These effects might be accounted for by the presence of an unusual poly(A) structure that protects the 5’-most adenosines from chemical modification (c) or an anomalous behavior of the reverse transcriptase in which the enzyme bypasses chemically modified adenosines after polymerizing through 6 modified adenosines (d).

These observations have potentially significant implications for current efforts to understand RNA structures and chemical modifications. Long stretches of adenosines appear throughout bacterial and eukaryotic messenger RNA, especially at their termini,^14-15^ and occur in numerous viral sequences, including the end of the HIV genome.^16^ Studies that seek to engineer unpaired loops into RNAs typically also use poly(A).^12, 17-18^ Nevertheless, understanding implications of the anomalous signal requires knowing the mechanistic origin of the poly(A) signature, especially if it is due to a special RNA structure in poly(A) stretches, as was previously hypothesized^12^; or to anomalies in how RTases respond to poly(A), analogous to effects seen in poly(A) processing by other molecular machines like the ribosome,^19^ RNA polymerase,^20^ or the RNAse H domains of HIV RTases.^21^ Here, we use alternative RNA substrates, capillary electrophoresis, nuclear magnetic resonance spectroscopy, and high-throughput sequencing to understand this mechanism. Our data disfavor the presence of an unusual structure in poly(A) RNA as an explanation of the anomalous signal. Instead, the results implicate an unexpected ability of RTases to bypass chemically modified adenosine after they have already polymerized through stretches of 4-6 adenosines, potentially triggered by an anomalous structure of poly(rA-dT) hybrids in the RTase active site.

## Results

### Generality of anomalous poly(A) signal across prior data sets

The suppression of apparent chemical reactivity in poly(A) was initially reported in ref.^12^ based on RMDB-deposited studies measuring SHAPE and DMS modification with the MAP-seq (multiplexed adduct probing read out by sequencing) method.^22^ To test that the anomaly was not an artefact of a single study or method, we compared data sets from numerous independent studies that used a variety of RTases and sequencing techniques (Fig. 2a). We catalogued each adenosine stretch in these sequences, and compared reactivities of adenosines that were at analogous positions relative to the end of each stretch, indexed as −1 (last adenosine in stretch), – 2 (second to last adenosine in stretch), etc. As control comparisons, we also compared mean reactivities of adenosines by their position relative to the beginning of each stretch (+1, +2, … positions; Fig. 2b). The RMDB MAP-seq data only show the suppression of chemical reactivity for adenosines at least 6 nucleotides 5’ to the end of poly(A) stretches (−7, −8, etc., in Fig. 2c; compare to +7, +8, etc., in 2d).

**Fig. 2.**
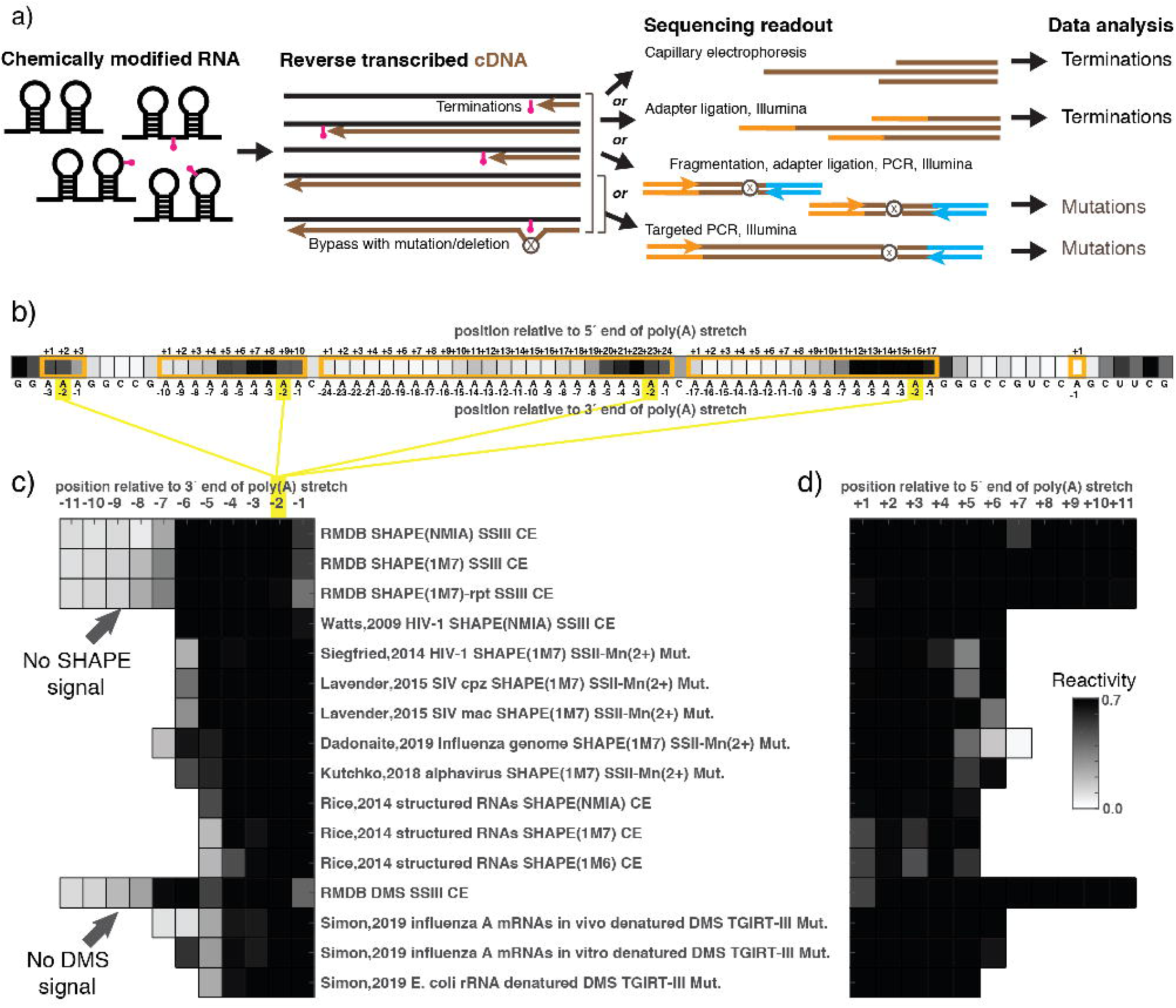
Survey of publicly available chemical mapping data support poly(A) anomaly. (a) Protocols for mapping RNA chemical modifications involves several steps, each with multiple possible variations. (b) Schematic for how adenosines within the single sequence shown in Fig. 1b are indexed before averaging DMS reactivity for positions based on positions within each polyadenosine stretch. (c) Reactivity data averaged according to position relative to 3’ end (left) or 5’ end (right) for each instance of a polyadenosine stretch. Reactivities within each data set were normalized so that the mean reactivity is unity at positions with maximal mean reactivity (typically −1 or +1 positions). References: RMDB (ETERNA_R69_0001)^11-12^; Watts, 2009^25^; Siegfried, 2014^10^; Lavender, 2015^26^; Dadonaite, 2019^27^; Kutchko, 2018^28^; Rice, 2014^24^; Simon, 2019^31^. RMDB – RNA Mapping Database, SHAPE – selective 2’-hydroxyl acylation and primer extension, NMIA – N-methyl isatoic anhydride, 1M7 – 1-methyl-7-nitroisatoic anhydride, 1M7 – 1-methyl-6-nitroisatoic anhydride, DMS – dimethyl sulfate; SS – SuperScript (reverse transcriptase), SSII-Mn(2+) – SuperScript II reverse transcriptase carried out with Mn^2+^ to promote mutational bypass, TGIRT – thermostable group II intron reverse transcriptase, CE – capillary electrophoresis (reads out reverse transcription termination), Term. – reverse transcription termination read out by Illumina sequencing, Mut. – reverse transcription incorporation of mismatches or deletions read out by Illumina sequencing.

We first investigated SHAPE data sets, for which the most data are currently available. On one hand, transcriptome-wide SHAPE studies across millions of nucleotide positions^23^ remain sparse, giving zero counts at many positions, and do not yet have necessary signal-to-noise to check for the poly(A) signal. On the other hand, SHAPE studies focusing on specific RNAs, such as viral genomes (HIV and SIV, alphavirus, and influenza genomes; around 10,000 nucleotides each) or collections of structured RNAs^24^, have higher signal-to-noise.^10, 25-28^ In several of these data sets, we detected suppression of SHAPE signal at −6 and −5 positions of polyadenosine stretches (labeled ‘Siegfried,2014’, ‘Lavender,2015’, ‘Dadonaite,2019’, and ‘Rice,2014’ in Fig. 2c). However, the data were sparse and control comparisons (Fig. 2c) suggested that the apparent protections might be explained by other effects such as base pairing of long adenosine stretches or biases in sequencing.^29^ Furthermore, there were few or no naturally occurring 7- or 8-adenosine stretches across these viral data sets, which would allow direct comparison to the MAP-seq SHAPE data that only show suppressions at −7, −8, etc. (arrow in Fig. 2c).

Beyond SHAPE, there is a growing body of publicly available DMS data on natural RNAs,^9, 30^ although nearly all that we checked were too poor in signal-to-noise to confirm or falsify the poly(A) anomaly. Fortunately, a recent study acquired DMS profiles for influenza A mRNAs and *E. coli* ribosomal RNA with excellent signal-to-noise.^31^ These data showed suppression of DMS signal at adenosines positioned at −5 within polyadenosine stretches (labeled ‘Simon, 2019’ in Fig. 2c), as did the original RMDB data studied in ref.^12^ (Fig. 2c), and the control comparisons show no analogous suppression at +1, +2, … to +6 positions (Fig. 2d). Furthermore, this study carried out DMS modification under denaturing conditions. These data therefore hinted that the suppression of the DMS signal might not be due to RNA base pairing, which would be disrupted in those experiments (Fig. 1c), but to later steps in the readout of the chemical modifications, as schematized in Fig. 1d and further tested below. These data also enabled a comparison of DMS signal across nearly all 5-mer sequences, which confirmed a striking specificity for a drop in DMS signal at position −5 in AAAAA and not in other purine-rich 5-mers (Fig. S1).

In these comparisons across different published data sets, differences in which adenosine positions showed apparent suppression in the different studies were complicated by the use of different RTases across the studies; different conditions of chemical modification including, in some cases, which reagent was used; differences in whether RTase termination or mismatch-introducing bypass was used to infer chemical modification; and differences in whether capillary electrophoresis or Illumina sequencing was used in the studies (Fig. 2a). To more directly confirm the anomalous signal, to dissect its origins, and to characterize the influence of different RTases, protocols, and readouts, we therefore turned to designed model systems.

### Anomalous poly(A) chemical mapping signal is recovered with simplest possible readout

As is apparent from the cross-study analysis of Fig. 2, dissecting the mechanism of the anomalous poly(A) chemical mapping signal is complicated by the large number of steps required in current structure mapping methods. The loss of signal at A’s could occur at any (or several) of these steps. For example, focusing on the chemical modification step, a single-helix structure of poly(A) with a period of six that leaves its 3’-most turn accessible to chemical modification could explain the observed SHAPE and DMS signals (see Fig. 1c and ref.^12^). Other steps with potential bias include the synthesis of the RNA by *in vitro* transcription with T7 RNA polymerase, which is known to ‘slip’ on repeated sequences^20^; similar slippage or biased termination of reverse transcriptases in the RTase step after chemical modification^32-33^; the ligation of adapters onto the resulting cDNAs, which are known to have sequence biases^22, 30, 34^; Illumina sequencing-by-synthesis of cDNAs used, which is also known to have sequence biases^29^; or the bioinformatic workup of these sequencing data into inferred chemical modification profiles.^29, 35^

We first sought to test if the poly(A) SHAPE and DMS signatures were due to the many possible biases occurring any steps downstream of reverse transcription, by carrying out the simplest method of reading out the cDNA products, capillary electrophoresis with fluorescently labeled primers. This readout detects RTase termination as shorter cDNA products. As a model sequence, we focused on an RNA developed in the Eterna project^12, 36^ called the Triangle of Doom-Sequence 1 (TOD-S1) (Fig. 3). TOD-S1 includes three stretches of 11 A’s each: two stretches are designed to be within the loops of hairpins jutting out of a three-way junction, and the third stretch is designed to be part of a single-stranded region near the 3’ end of the construct. Flanking hairpins provide normalization standards (Fig. 3a).^37^ Unlike the MAP-seq protocol, where RNAs were synthesized in a large pool with other molecules, these RNAs were synthesized individually by T7 RNA polymerase from synthetic DNA templates. After chemical modification and reverse transcription, we directly visualized the cDNA products through capillary electrophoresis (CE), instead of ligating on adapters and carrying out Illumina next-generation sequencing.

**Fig. 3.**
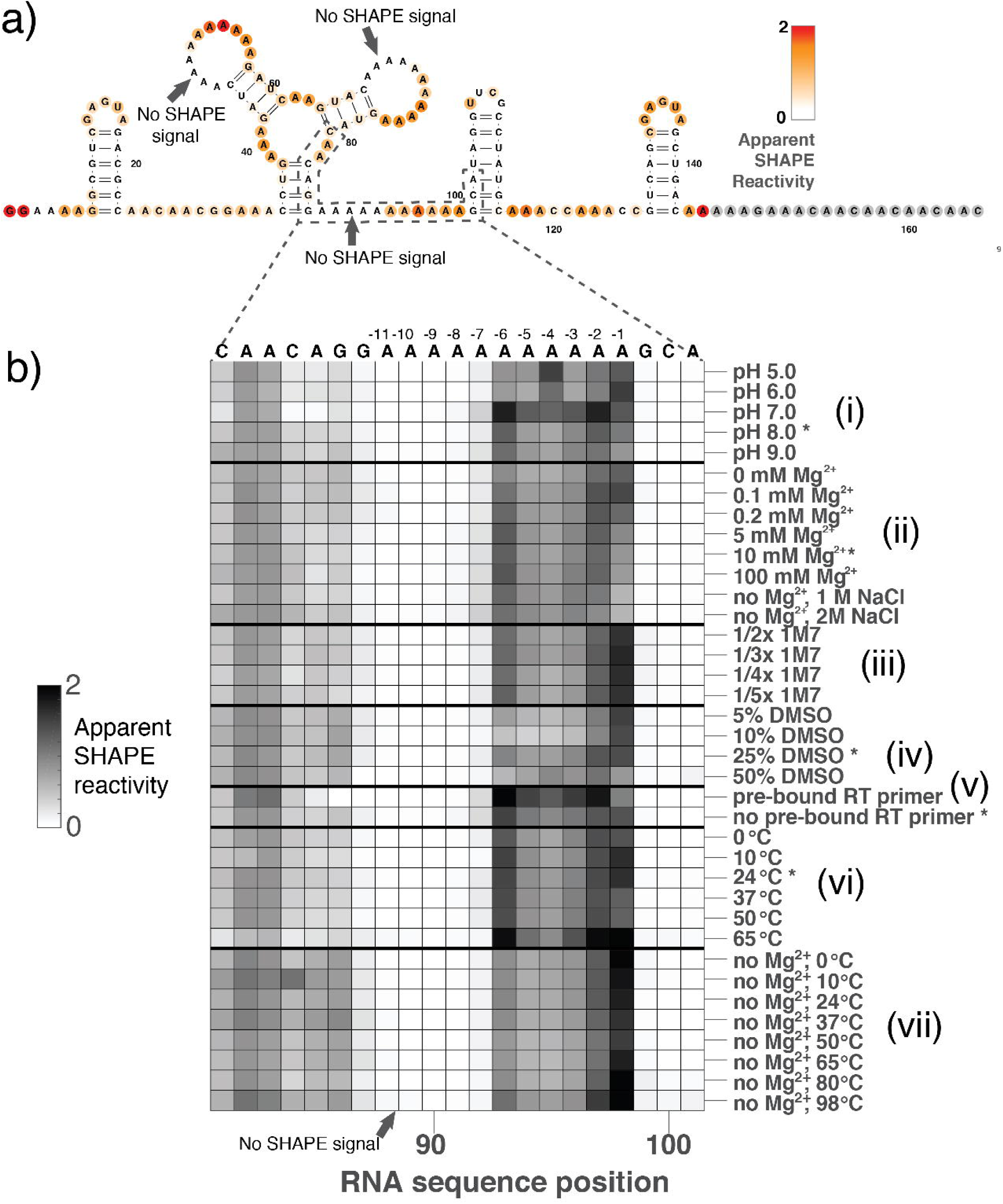
Anomalous SHAPE signals in the TOD-S1 model RNA. (a) Apparent SHAPE reactivity of the TOD-S1 construct, which contains three stretches of 11 adenosines. (b) Anomalous SHAPE profile is robust to a wide variety of changes to solution conditions under which TOD-S1 RNA was chemically modified. Data for the most clearly resolved (3’-most) A11 stretch of the TOD-S1 construct are shown. Standard solution conditions for chemical modification were 10 mM MgCl_2_, 50 mM Na-HEPES, pH 8.0 at ambient temperature (24 °C), with cosolvent of 25% anhydrous DMSO used to the SHAPE reagent 4.24 mg/ml 1M7. Perturbations to these conditions are labeled on the *y* axis of (c), with asterisks (*) marking replicates of the standard conditions. Data shown have been background subtracted based on control reactions without SHAPE modification reagent, normalized so that 1.0 corresponds to the average reactivity of GAGUA hairpin loops included at 5’ and 3’ ends of the RNA as internal controls. Anomalous regions with no SHAPE signal highlighted with gray arrows.

Fig. 3a shows the CE-based SHAPE data as coloring on the TOD-S1 secondary structure. For each stretch of 11 A’s, there was clear evidence for strong SHAPE modification for the last 6 A’s, but few or no detectable cDNA products corresponding to modification of the first 5 A’s of the same stretch. As above, we have numbered positions in a poly(A) stretch relative to the 3’ end; so here only positions −6 through −1 appeared SHAPE-modified, whereas positions −11 to −7 did not show terminated cDNA products corresponding to SHAPE modification. Similar SHAPE patterns were visible for the 5’-most and middle stretch of A’s in these capillary electrophoresis data for TOD-S1. These patterns corresponded well to analogous measurements in the RMDB MAP-seq experiments.^12^ Further CE-based data involving constructs with C mutations in the TOD-S1 loops (called TOD-S7), and using DMS instead of SHAPE, further matched the observations in the prior MAP-seq data (Fig. S2). These CE data demonstrated that the anomalous poly(A) signatures observed in MAP-seq and other literature data sets (Fig. 2c) were not due to biases in the Illumina sequencing workup of cDNAs since those workup steps are absent in this capillary electrophoresis readout.

### Alternative solution conditions during chemical modification give the same poly(A) anomaly

The lack of truncated cDNA’s corresponding to SHAPE or DMS modification at the middle of the poly(A)’s could be due to an unusual conformation at those positions that might sterically protect the 2’-hydroxyl or N1 position of the adenosine from chemical modification (Fig. 1c). We reasoned that such a poly(A) structure may be destabilized or at least modulated by solution conditions, and so we carried out six sets of experiments to test this prediction (Fig. 3b).

At high RNA concentrations and low pH, poly(A) strands are known to form a parallel double helix, which is stabilized by protonation at the adenosine N1 position.^38^ We varied the pH of our measurements away from our standard pH of 8.0 from 5.0 to 10.0; however we saw no change in the poly(A) SHAPE profiles of TOD-S1 (Fig. 3b-(i)). Second, RNA secondary and tertiary structures are generally stabilized by the presence of Mg^2+^, which was present at a concentration of 10 mM MgCl_2_ in all above measurements, and the stability can be modulated by high concentrations of monovalent salt. However, when we repeated our measurements without Mg^2+^ or with lower concentrations of Mg^2+^, or with 1 M or 2 M NaCl (and no Mg^2+^), we again saw no change in SHAPE profiles of TOD-S1 (Fig. 3b-(ii)) Third, we asked if the modification reagent or its cosolvent (anhydrous DMSO) might be stabilizing an alternative poly(A) structure, so we varied the concentration of the SHAPE modifier 1M7, going as low as one fifth of our standard concentration, but saw similar patterns (Fig. 3b-(iii)). Fourth, we directly added the SHAPE cosolvent DMSO to up to 50% concentration, and again saw no change in the poly(A) regions, though we observed perturbed chemical modification profiles outside those regions confirming the denaturing effect of DMSO (Fig. 3d-(iv); Fig. S2a,b). Fifth, our constructs for MaP-seq and the current CE-based measurements all include an A-rich primer binding site at the 3’ end of the sequence (gray nucleotides, Figs. 1a-b, 3a). To test if this region might be interacting with poly(A) sequence, we repeated measurements with the reverse transcription primer present during chemical modification as a competitive inhibitor of a possible interaction; however we observed the same pattern as before (Fig. 3b-(v)). Sixth, we reasoned that if poly(A) forms an unusual structure, we would be able to modulate its stability relative to an unfolded configuration by increasing the temperature. However, in SHAPE measurements ranging from 0 °C to 98 °C, we observed no changes in the poly(A) regions, either with or without MgCl_2_ (Fig. 3b-(vi) and Fig. 3b-(vii)). Data from these six sets of experiments strongly disfavored a model where the observed SHAPE signature at poly(A) might be due to an anomalous RNA structure. Further CE-based experiments with the TOD-S7 variant and using DMS instead of SHAPE gave additional support to these conclusions (Fig. S2a,b).

### NMR spectroscopy further disfavors the presence of an unusual poly(A) RNA structure

As a further test of whether poly(A) forms an anomalous structure, we synthesized model RNA molecules and carried out NMR spectroscopy under conditions matching our chemical mapping experiments (50 mM Na-HEPES pH 8.0, 10 mM MgCl_2_; 25 °C). For an RNA consisting of 12 adenosines, all of the resonances fell in the 2D [^13^C,^1^H] HSQC chemical shift region similar to results for a model RNA shown to adopt a stacked conformation that is in equilibrium with the unfolded state (compare red contours in Fig. 4a and 4b; ref.^39^). In the SHAPE and DMS chemical mapping experiments described above, substitution of individual A’s to C, G, or U produced large changes in apparent chemical reactivity for stretches of six adenosines 3’ to the substitution (Fig. 1). By NMR however, such substitutions gave only small changes in chemical shifts, and these changes were limited to adenosines immediately neighboring the substitution and not for longer stretches of 6 adenosines 3’ to the substitution (Fig. 4b,c). In addition, 2D NOESY spectra indicated that the nucleotides adopt a predominantly anti conformation with relatively weak C1’-C8 NOE cross peaks (Fig. S3). Thus, NMR experiments disfavor a noncanonical RNA structure as an explanation for the anomalous chemical mapping signals in polyadenosine, in agreement with the biochemical data above.

**Fig. 4.**
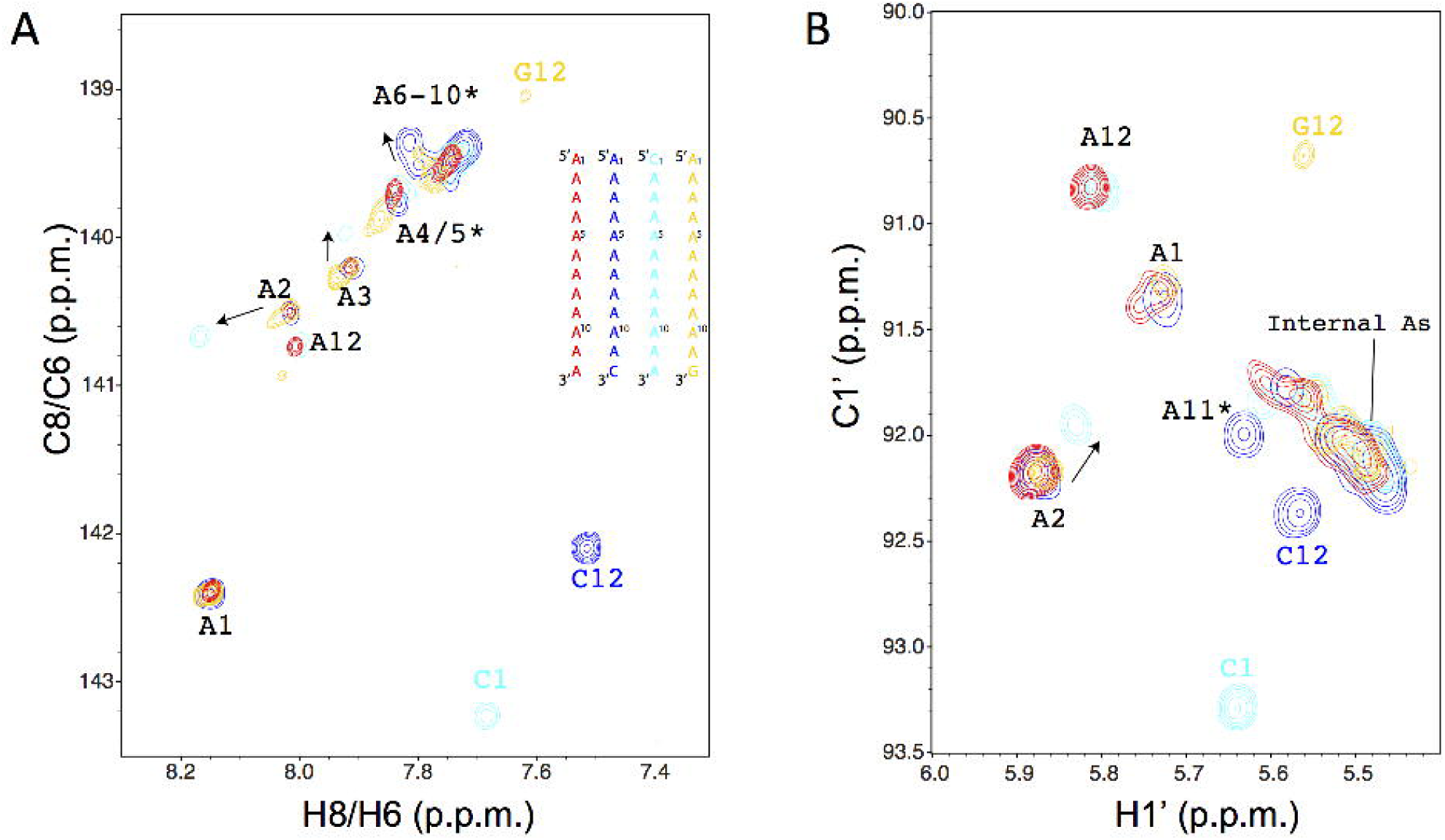
NMR spectra indicate A-form like conformations throughout polyadenosine. (a,b) Chemical shifts of C8 and H8 atoms in [^13^C,^1^H] HSQC experiments fall within the range of A-form helices for both a reference sequence previously shown to form a single-stranded stacked configuration^62^ (a) and for a 12-adenosine RNA A_12_ (b). In (b), substitutions of the 5’ adenosine of A_12_ to cytidine (CA_11_, cyan) or the 3’ adenosine to cytidine (A_11_C, blue) or guanosine (A_11_G, gold) give small perturbations to chemical shifts of immediately neighboring bases (arrows) but no evidence for dramatic perturbations or induction of conformational rearrangements extending six nucleotides. (c) Chemical shifts in the aromatic base region from the same [^13^C,^1^H] HSQC experiments also appear in the range of conventional A-form helices. Spectra acquired in Na-HEPES buffer (1 mM RNA in 50 mM Na-HEPES pH 8.0, 10 mM MgCl_2_) at 25 °C.

### Alternative reverse transcriptases give distinct poly(A) signatures

Given the lack of evidence for adapter ligation or Illumina sequencing biases (Fig. 3) or an anomalous poly(A) structure (Fig. 3-4) as an origin of the SHAPE/DMS anomaly, we developed experiments to instead test for an unexpected behavior in the intermediate step of reverse transcription (Fig. 2a). We considered a model in which the RTase bypasses chemically modified adenosines that are positioned at least six nucleotides ahead of the end of a poly(A) stretch (Fig. 1d). In this model, the RTase does not terminate or make a mutation, but instead skips by correctly polymerizing a dT or several dT’s (insertion) as it bypasses the chemically modified A; any or all such events would lead to absence of a termination product detected by capillary electrophoresis and explain the anomalies of Figs. 1-3.

One prediction from this RTase bypass model is that different reverse transcriptases could give different cDNA profiles when reverse transcribing the same pools of SHAPE-modified or DMS-modified RNAs. We therefore tested the Moloney Murine Leukemia Virus (MMLV) reverse transcriptase and its commercially available variants SuperScript II, III, and IV. On one hand, MMLV and SuperScript II both gave similar patterns to our original standard RTase SuperScript III, giving cDNA termination products at positions −6, −5, to −1 for SHAPE-modified RNA (Fig. 5, top). In contrast, SuperScript IV gave cDNA termination products only at two positions −2 and −1, i.e., the last two adenosines of each poly(A) stretch, and efficiently bypassed modifications at more 3’ positions. Interestingly, the behavior of SuperScript IV was nearly identical to the behavior of SuperScript II reverse transcribing in the presence of Mn^2+^, conditions recommended for mutational bypass.^10^ This observation suggests that the MMLV-derived enzymes have an alternative ‘mode’ of bypass that can be triggered either by Mn^2+^ or by mutations installed into SuperScript III to create SuperScript IV. Going beyond the MMLV family, we further tested two RTases derived from thermostable group II introns, TGIRT-III^9, 34^ and MarathonRT^40^. Both of these enzymes also nearly completely bypassed SHAPE modification and DMS modifications in poly(A) stretches (Fig. 5, bottom). Additional experiments with the TOD-S7 variant confirmed these conclusions (Fig. S2c).

**Fig. 5.**
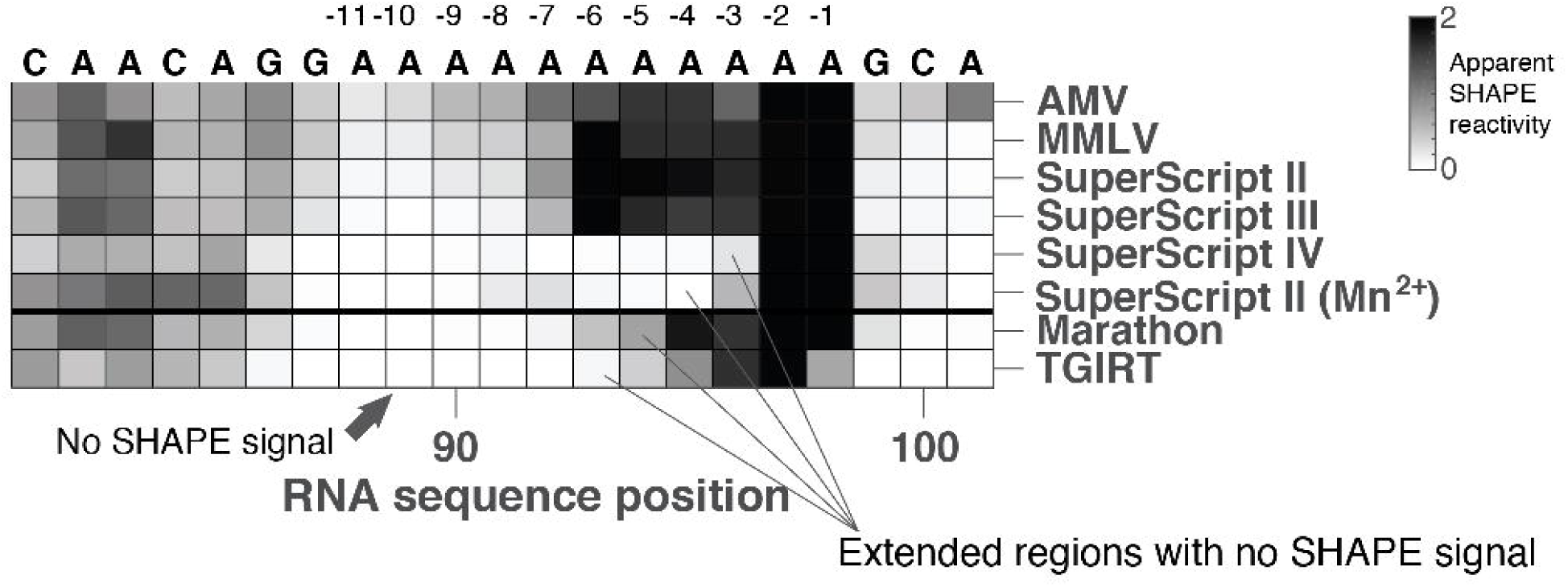
Anomalous SHAPE signals change with reverse transcriptase used to read out chemical modifications. Data for the most clearly resolved (3’-most) A11 stretch of the TOD-S1 construct are shown. AMV – avian myeloblastosis virus, MMLV – Moloney murine leukimia virus, TGIRT – thermostable group II intron reverse transcriptase (variant TGIRT-III).

To test readouts different from capillary electrophoresis (Fig. 2a), we used adapter ligation followed by Illumina sequencing and discovered analogous differences in termination and in mismatch-free bypass in poly(A) depending on which RTase was used (Fig. S4), further implicating the RTase as a likely culprit for the poly(A) anomaly. As a final series of SHAPE-based tests, we designed new constructs that introduced stretches of U’s that were distal to the poly(A) stretches of TOD-S1 but base paired to these originally single-stranded poly(A) stretches. The resulting perturbed structures were predicted to give distinct SHAPE patterns in the anomalous structure vs. RTase bypass models (Fig. S5a-d). As shown in Fig. S5e, the actual experimental results strongly favored the RTase bypass model and disfavored the anomalous structure model, in agreement with our previously described analyses (Figs. 1-5).

### Near-quantitative bypass at m^1^A modifications installed during chemical synthesis

All of the structure mapping results above from prior studies and our newer experiments are consistent with near-quantitative RTase bypass within poly(A) stretches. However, these experiments relied on a relatively uncontrolled step: installation of chemical modifications at random positions throughout the RNA molecules via SHAPE or DMS treatment. To remove this this randomness, we sought to install chemical modifications specifically at each position of a poly(A) stretch. The RTase bypass model makes predictions for how the RTase should terminate or bypass chemical modifications at each position. For example, DMS modification results in m^1^A nucleotides. Our DMS results above would therefore predict that SuperScript III RTase should terminate at m^1^A’s installed at the −6, −5, −4, −3, −2 to −1 positions, with particularly strong termination at −3 and −2 and weakest termination at −5 (Fig. 2c). The RTase should bypass m^1^A’s installed at any positions further 5’ of these last 6 adenosines (Figs. 1-3). We tested these predictions on chemically synthesized substrates with m^1^A at each position in an 11-adenosine stretch (Fig. 6). For these experiments, we shifted to a smaller 41-nucleotide RNA construct called Syn41 to ensure reasonable synthesis yields, and confirmed that chemically synthesized Syn41 gave a poly(A) DMS and SHAPE pattern similar to our previous enzymatically synthesized substrates (Fig. 6a).

**Fig. 6.**
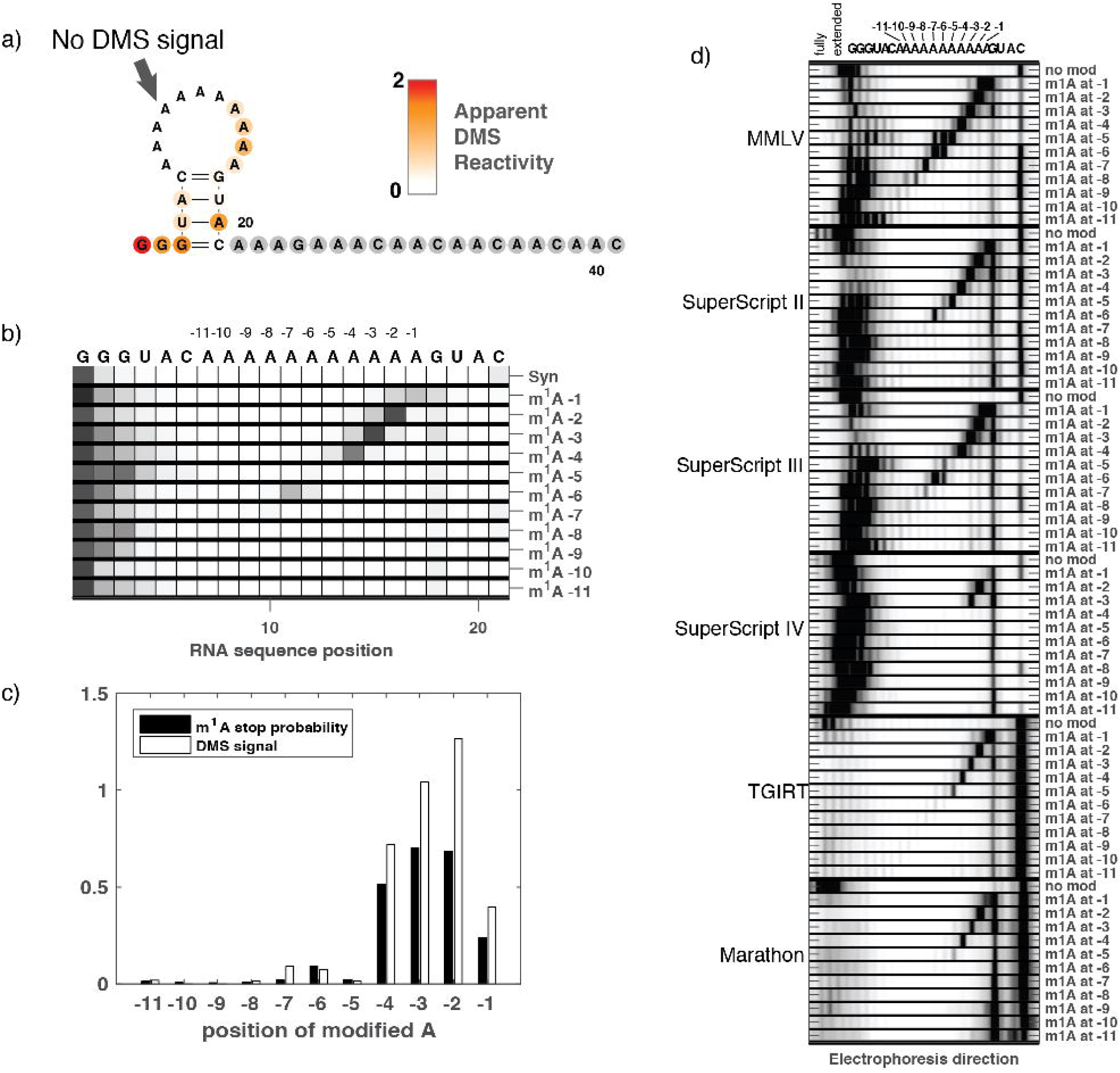
Reverse transcriptase read-through at 1-methyl-adenosines incorporated into a poly(A) stretch during synthesis. (a) DMS mapping data on the Syn41 construct. (b) Termination probabilities of SuperScript III reverse transcriptase on constructs that each present a single m^1^A installed at the positions specified on *y* axis. Note that modifications at −11 through −8 do not terminate the reverse transcriptase. (c) The measured termination probabilities explain the anomalous signal seen in dimethyl sulfate probing experiments read out by SuperScript III, where no signal is apparent at −11 to −8, and the maximum signal appears at −4 to −2. (e) Electropherograms of cDNA extended from Syn41 that is not modified (no mod) or m^1^A-modified, as reverse transcribed from a large range of RTases. Sequence assignment of bands is tentative for longer products (towards left of electropherograms), which may contain insertions and deletions.

The reverse transcription results across m^1^A-harboring RNAs are shown in Fig. 6b. The data agree well with the predictions of the RTase bypass model. SuperScript III terminated nearly quantitatively at m^1^A installed at the −3 and −2 positions, with very little full-length cDNA product appearing. The enzyme partially terminated and partially read through m^1^A at the other six 3’-most positions of the 11-adenosine stretch. Strikingly, SuperScript III bypassed m^1^A installed at −11, −10, −9, −8, and −7 positions with near-quantitative efficiency, resulting in near-full-length cDNAs (Fig. 6d). This pattern of bypass efficiencies was in excellent agreement with the DMS patterns at poly(A) stretches measured in earlier experiments (Fig. 6c). Experiments with other reverse transcriptases on these m^1^A substrates also gave bypass efficiencies that accorded with DMS patterns measured previously in other laboratories or by us (Figs. 2, 4 to 6d). For example, the group II intron derived RTases (TGIRT-III and MarathonRT) efficiently reads through m^1^A at all positions except −4, −3, −2, and −1 (Fig. 6d), similar to their behaviors in literature (label ‘Simon,2019’ in Fig. 2) and our CE-based (Fig. S2c) DMS experiments. These comparisons gave strong support for the RTase bypass model, based on prospective experiments.

These experiments also gave new information on how the RTase might bypass chemical modification. Deletions, incorporation, or insertions across from m^1^A would give rise to extended cDNA products shorter, the same size, or longer than products reverse transcribed from unmodified RNA. For the MMLV/Superscript family of reverse transcriptases (Fig. 6d, left), we observed products that were 1-5 nucleotide shorter in m^1^A -containing substrates compared to unmodified substrates, suggesting that these enzymes bypass m^1^A with deletions. Interestingly, the distribution of cDNA lengths depended on the position of the chemical modification in the poly(A) stretch, with m^1^A at –5 giving not only shorter bypass products but also a spread in termination products (see, in particular, MMLV, Fig. 6b) that extend 1 or 2 nts beyond the m^1^A rather than cleanly terminating at the modification site. Intriguingly, the group II intron derived RTases (TGIRT-III and MarathonRT) showed near-quantitative bypass of m^1^A installed at most positions, but the extended products appeared as fainter bands spread across a wider and longer distribution of lengths than the MMLV/Superscript RTases (Fig. 6b). The different distributions of bypass products indicate that the molecular details of bypass are different between enzymes.

### Implications for RNA structure inference

The observation that RTases can bypass modifications in stretches of adenosines raises concerns for chemical mapping experiments used to biochemically probe RNA structure. After carrying out the studies above, we re-examined data we had been acquiring on the HIV 3’ untranslated region (3’ UTR). This viral RNA segment is polyadenylated during its biogenesis *in vivo* and is hypothesized to exhibit functionally important structures which may interconvert during different viral stages and recruit distinct partners.^10, 16, 41^ We had originally seen paradoxical results when carrying out DMS mutational profiling^9^ and mutate-and-map-seq experiments^42^ without and with a poly(A) tail. On the one hand, we observed almost no difference across the entire 3’ UTR’s DMS-MaP-seq profile in constructs without and with the poly(A) tail (Fig. 7a). Here, the poly(A) tail was represented by a 20-adenosine stretch placed after the 3’ UTR polyadenylation site (the total stretch length is 24 adenosines due to an additional A at the polyadenylation site and three A’s from our appended primer binding site). The DMS profile comparison suggested that there is negligible structural effect from polyadenylation, at least for RNA probed *in vitro*. However, structure modeling suggested a different picture. Modeling based on RNAstructure’s *Fold* executable^43^ guided by the DMS and mutate-and-map-seq data^42^ (Fig. S6) resulted in the secondary structures shown in Figs. 7b and 7c. In the structure model without the poly(A) tail, stems corresponding to three USE (upstream sequence domain) hairpins and the long TAR (trans-activation response) element were modeled with high confidence (based on bootstrap uncertainty estimation^42, 44-45^; see inset of Fig. 7b and Fig. S6). However, inclusion of the poly(A) sequence resulted in a rearrangement in which one of the USE hairpins (sequence U_5_, blue in Fig. 7) shifted to base pair with five of the introduced A’s (yellow, Fig. 7b), in contradiction with the near indistinguishability of the DMS profile at and around that region (Fig. 7a).

**Fig. 7.**
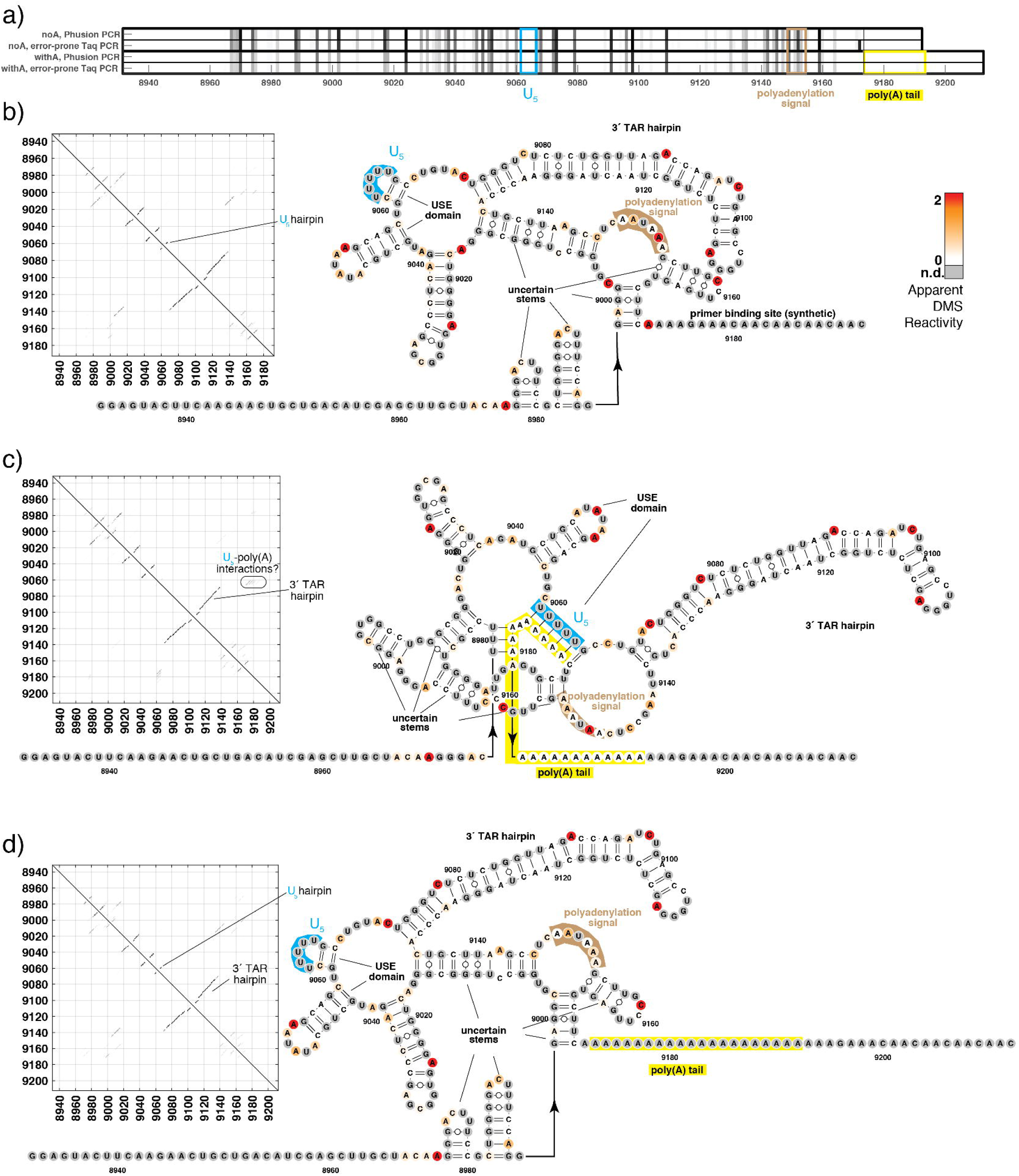
Reverse transcriptase read-through of DMS-modified nucleobases leads to anomalous reactivity data that can confound structural modeling of the HIV 3’ UTR by chemical mapping. (a) DMS profiles of without and with A_20_ tail are nearly indistinguishable. Reverse transcription through the poly(A) tail of the HIV 3’ UTR demonstrates a striking loss of apparent DMS reactivity (white). (b-d) Predicted secondary structures of HIV 3’ UTR without A_20_ tail (b), with A_20_ tail and using the conventional assumption that DMS protections in the tail reflect formation of structure (c), and with A_20_ tail but ignoring DMS protections (gray) due to invisibility of chemical modifications to the RTase readout (SuperScript II with Mn^2+^).

The contradiction is resolved by the findings described above that chemical modifications within poly(A) are essentially invisible to reverse transcription. In accord with this picture, we observed no apparent DMS reactivity in the poly(A) tail appended to the HIV 3’ UTR. Any mismatches in the last three A’s were masked by use of a reverse transcription primer including those A’s, and we expected the RTase (SuperScript II with Mn^2+^) would bypass any chemical modifications at the 21 previous A’s and therefore give no DMS signal there (Fig. 5). Given this bypass, the correct use of *Fold* is to not apply any energetic bonuses or penalties in the poly(A) stretch, analogous to treatment of G and U residues that do not give DMS signals in our conditions (gray nucleotides in Fig. 7). Ignoring poly(A) data gives the model structure shown in Fig. 7d. The poly(A) is modeled as fully unpaired, and this secondary structure recovers all the base pairs of the poly(A)-less HIV 3’ UTR without any rearrangements. These results illustrate the importance of taking into account anomalous reverse transcription at poly(A) when interpreting chemical mapping data to infer RNA structures.

## Discussion

DMS and SHAPE mapping experiments give anomalous signatures at poly(A) stretches: a striking loss of reverse transcription cDNA termination products for adenosines that are at least six nucleotides upstream of a poly(A) 3’ end. Our results and data from previous studies show that a single nucleotide substitution in the poly(A) stretch abrogates this anomalous behavior. We have dissected each step of our structure mapping protocol to understand the mechanism of anomalous poly(A) DMS and SHAPE signatures. Experiments that attempted to ‘melt’ a possible structure of poly(A) RNA or to displace it through base pairing failed to confirm any anomalous structure; and NMR experiments also disfavored such an anomalous structure. Instead, the DMS and SHAPE signatures varied with the reverse transcriptases used to read the signatures, with some RTase enzymes showing nearly quantitative bypass of termination after reverse transcribing six adenosines and others shifting to bypass chemical modifications with fewer adenosines. Analogous measurements on RNA substrates with m^1^A installed at specific sites during chemical synthesis confirm that RTases bypass instead of terminate at the modifications. The efficiency of bypass depends sharply on the number of A’s that the RTase has already polymerized and on which RTase is used. Some recently engineered RTases (SuperScript IV, TGIRT-III, MarathonRT) bypassing chemical modifications after polymerizing as few as 2-3 adenosines.

The behavior we observe with long polyadenosine stretches is compatible with available knowledge of the enzymatic activities and structures of reverse transcriptases. For example, the series of SuperScript enzymes developed from MMLV have a terminal nucleotidyl transferase (TdT) activity that favors addition of dT to cDNA/RNA hybrids^46^, although this activity has been suppressed through engineered mutations in the SuperScript enzymes. Classic studies on the HIV-1 RTase at poly(A) stretches revealed template-primer slippage of RTase, resulting in insertions or deletions^32, 47^ or template switching.^48^ It is not yet clear if the near-quantitative bypass mode we observe, which is triggered by a specific length of poly(A) (six nucleotides for SuperScript III), involves the same molecular steps as the TdT activity or the slippage mechanisms previously described. In particular, the previous experiments did not test whether TdT or the observed slippage events have a sharp and extended dependence on poly(A) length or the same sensitivity to even single-nucleotide changes like A to U. Such studies will be necessary to understand the connections with the various enzymatic activities that have been previously uncovered and the bypass described here. Other papers have noted a dependence of reverse transcription on nucleotides immediately 3’ to the site of polymerization across a variety of RTases^7, 49-52^ but again have not systematically varied the lengths of the poly(A) stretches. Further experiments will be necessary to understand these potentially complex and sharp dependences on 3’ sequence and length and on RTase type.

From a structural point of view, a dependence of RTase activity on a previously reverse-transcribed stretch of adenosines is plausible. As illustrated in Fig. 8a, an RTase paused at a chemical modification would be sensitive to fluctuating structures of a previously polymerized rA/dT hybrid. Such structural fluctuations might enable bypass through the roadblock, e.g., by repositioning of the RNA template to bring the next (unmodified) ribonucleotide into the active site. Consistent with this model, crystallographic studies of the HIV-1 reverse transcriptase show that the cDNA/RNA hybrid is held within the RTase for numerous base pairs beyond the site of cDNA polymerization (Figs. 8b), and the RTase active site domains are well-positioned to engage with the sequence of the RNA/cDNA hybrid that is polymerized immediately prior to the RNA in the active site. Other RTase domains such as the RNAse H domain are more distal from the polymerization active site but can indirectly readout changes in the shape of the hybrid near the active site. Indeed, biochemical studies have shown that contacts of the cDNA/RNA hybrid with the HIV-1 RTase enzyme’s RNAse H domain (∼18 bp away from the polymerase active site) can be modulated by changes in the hybrid sequence.^53-54^ Furthermore, crystallographic studies have revealed that the docking mode of the cDNA/RNA hybrid within the RTase changes with the hybrid sequence (compare sub-panels of Fig. 8b). ^21 51^ In future mechanistic work, bringing a long (greater than 6 bp) A/dT hybrid into the polymerase active site may help stabilize the ‘bypass’ mode of reverse transcriptases and enable incisive kinetic and structural studies by crystallography or cryo-electron microscopy (Fig. 8b, bottom-right sub-panel).

**Fig. 8.**
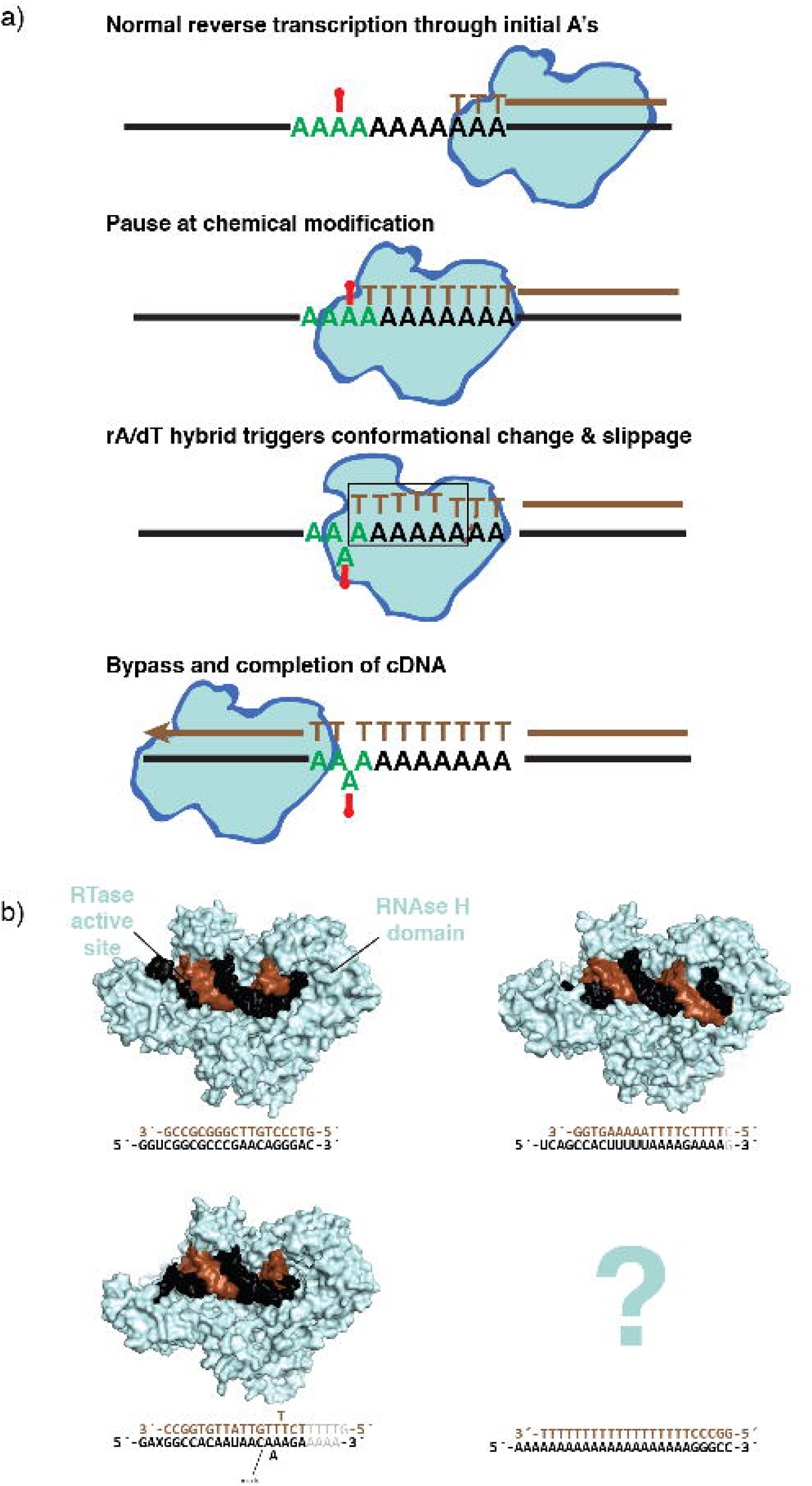
Possible mechanism for anomalous reverse transcription bypass at chemically modified poly(A) substrates. (a) Reverse transcriptase enzyme (cyan) polymerizing DNA (brown) on an RNA template (black) bypasses chemical modifications at which it would normally terminate (red) if it has previously reverse transcribed adenosines. The mechanism may involve recognition of the poly A-dT hybrid duplex (green) by the enzyme. (b) RNA/DNA hybrids of different sequences show different docking modes to the HIV-1 reverse transcriptase (from bottom-left, clockwise, PDB IDs: 6BSI^21^, 4PQU^63^, 1HYS^53^). To aid visual comparison of differences, only the first 18 RNA and DNA nucleotides 3’ and 5’ to the active site, respectively, and any 5’ RNA overhangs are shown in the 3D models. Other base pairs denoted in gray at the sequence depiction at bottom of each panel. RNA – black, DNA – brown, cyan – reverse transcriptase.

The biological significance of bypass of poly(A) chemical modifications is not yet clear, but retroviral RNA genomes and their functionally important cleavage products are polyadenylated; the genomes also harbor functionally important polypurine stretches (although these tracts have both G and A).^53-55^ We speculate that RTase bypass of chemical modifications at these sites may be important for the retrovirus to evade host responses involving chemical modification of viral RNAs. Our study has highlighted the same anomalous bypass effect in group II-intron-derived, rather than retroviral, reverse transcriptases. It is unclear whether these enzymes, which are also parts of selfish elements^56^, also encounter chemically modified poly(A) sequences during their evolution.

In addition to biological implications, our study has implications for two classes of biotechnology methods that involve detection of RNA chemical modifications. First, given our results on the HIV 3’ UTR DMS mapping and structural inference, 3’ sequence context effects may be introducing systematic biases in structural studies based on chemical mapping using, e.g., the SHAPE-MaP and DMS-MaP-seq approaches. Second, 3’ sequence context effects may be causing m^1^A and chemical modifications in brain mRNA and other natural samples to be bypassed by RTases and therefore undercounted, perhaps severely so in poly(A) tails. For both applications of reverse transcriptases, correcting these biases will require detailed characterization of sequence contexts of reverse transcription termination vs. read-through, extending up to six nucleotides 3’ of the modified nucleotides and specially performed for each RTase enzyme used or newly developed.

## Supporting information

DNA and RNA Sequences

## Acknowledgments

We thank Eterna participants J. Anderson-Lee, E. Fisker, R. Wellington-Oguri, M. Wiley, and M. Zada, and S. Rouskin (MIT) for discussions; and K. Liu and J. Yesselman for help with pilot studies and analysis. We thank A. Pyle (Yale University) for the gift of the MarathonRT enzyme. This study was funded by the National Institutes of Health through grants R01 GM100953, R35 GM122579, R21 CA219847, R21 AI145647, and P50 GM 103297.

## Methods

### Enzymatic nucleic acid preparation and purification

RNA was transcribed using DNA templates containing a T7 RNA Polymerase promoter sequence at their 5’ ends and a 20 base-pair Tail2 sequence at their 3’ ends (Supplemental File, sequences.xlsx). The RNA sequence consisted of the sequence of interest flanked by a reference hairpin on each side serving as internal structural controls. The DNA template was assembled through DNA primer assembly using primers designed using the Primerize tool (https://primerize.stanford.edu/).^57^ Designed primers were ordered in plate format from Integrated DNA Technologies (IDT) and assembled via PCR assembly with Phusion DNA polymerase as described on the Primerize PCR Assembly Protocol (https://primerize.stanford.edu/protocol/#PCR). Assembly products were verified for size via agarose gel electrophoresis and subsequently purified using Agencourt RNAClean XP beads. Purified DNA was quantified via NanoDrop (Thermo Scientific) and 8 pmol of purified DNA was then used for in vitro transcription with T7 RNA polymerase (New England Biolabs Inc.). The resulting RNA was purified with Agencourt RNAClean XP beads supplemented with an additional 12% of PEG-8000 (3 volumes of 40% PEG-8000 was added to 7 volumes Agencourt RNAClean XP beads) and quantified via NanoDrop. The sequences of the transcribed Triangle of Doom (TOD) RNA molecules, including reference hairpins, and of transcribed stem-extended TOD-S1 and TOD-S7 RNA molecules, are provided in Supplemental File sequences.xlsx.

### RNA modification

1.2 pmol of RNA was denatured in 50 mM Na-HEPES pH 8.0 at 90 □ for 3 minutes and cooled at room temperature for 10 minutes. The RNA was then folded with the addition of MgCl_2_ to a final concentration of 10 mM in 15 μL and incubated at 50 □ for 30 minutes, then left at room temperature for 10 minutes. For chemical modification of folded RNA, fresh working stocks of dimethyl sulfate (DMS, Sigma-Aldrich) and 1-methyl-7-nitroisatoic anhydride (1M7) were prepared. For DMS, 1 μL of DMS was mixed with 99 μL of 100% EtOH. The 100 μL solution was then added to 100 μL RNase free H_2_O for a final volume of 200 μL. For 1M7, 4.24 mg of 1M7 was dissolved in 1 mL of anhydrous DMSO. In the modification step, 5 μL of modifying agent (either DMS or 1M7) working stock was added to 15 μL of folded RNA and incubated at room temperature for 15 minutes. Modification reactions were quenched with 5 μL of quenching solution (β-mercaptoethanol or 0.5 M Na-MES pH 6.0 for DMS and 1M7, respectively). Then, 5 μL of 5 M NaCl, 1.5 μL of oligo-dT Poly(A)Purist MAG beads (Ambion), and 0.065 pmol of 5’ Fluorescein (FAM)-labeled A20-Tail2 primer were added (see Supplemental File sequences.xlsx), and the solution was mixed and incubated for 15 minutes. The magnetic beads were then pulled down by placing the mixture on a 96-post magnetic stand, washed twice with 100 μL of 70% EtOH, and air dried for 10 minutes before being resuspended in 2.5 μL RNase-free H_2_O.

### cDNA synthesis

The 2.5 μL resuspension of purified, polyA magnetic beads carrying chemically modified RNA was mixed with 2.5 μL of reverse transcription mix with SuperScript-III (Thermo Fisher), or an alternate reverse transcriptase as otherwise specified (Table S1). The reaction was incubated at 48 □ for 60 minutes. The RNA was then degraded by adding 5 μL of 0.4 M NaOH and incubating the mixture at 90 □ for 3 minutes. The degradation reaction was placed on ice and quickly quenched by the addition of 2 μL of an acid quench solution (1.4 M NaCl, 0.6 M HCl, and 1.3 M NaOAc).

### Measurement of chemical reactivity with capillary electrophoresis

Bead-bound, FAM labeled cDNA was purified by magnetic bead separation, washed twice with 100 μL of 70% EtOH, and air-dried for 10 minutes. To elute the bound cDNA, the magnetic beads were resuspended in 10.0625 μL ROX/Hi-Di (0.0625 μL of ROX 350 ladder, Applied Biosystems, in 10 μL of Hi-Di formamide, Applied Biosystems) and incubated at room temperature for 20 minutes. The resulting eluate was loaded onto capillary electrophoresis sequencers (ABI-3100 or ABI-3730) either on a local machine or through capillary electrophoresis services rendered by ELIM Biopharmaceuticals. Data were analyzed, background-subtracted, and normalized with HiTRACE.^37, 58^

### Varying chemical mapping experimental conditions

For both the TOD-S1 and S7 RNA constructs, the chemical mapping experiments described above were carried out under a variety of conditions. The pH of the RNA denaturation and folding steps was varied from 5.0 through 10.0 by using 50 mM of one of the following folding buffers: Na-MES pH 5.0, Na-MES pH 6.0, Tris-HCl pH 7.0, Na-HEPES pH 8.0, Na-CHES pH 9.0 and Na-CHES pH 10.0. For 1M7, final working modifier concentrations of 0.1, 0.52, 0.24, and 1.06 mg/mL in DMSO were tested. For DMS, final modification reaction concentrations of 0.25%, 0.125%, 0.06%, and 0.03% were tested. For folding reaction salt concentrations, 0, 0.1, 0.2, 5.0, 10.0, and 100 mM of MgCl_2,_ or 1.0 and 2.0 M of NaCl (with no MgCl_2_) were tested. Final concentrations of DMSO in each working modifier stock were tested at 0%, 5%, 10%, 25%, and 50% for DMS; 5%, 10%, 25%, 50% for 1M7. The folding and modification steps were tested with and without the presence of 0.6 pmol of the FAM-labeled A20-Tail2 primer. Modification reaction temperatures of 0, 10, 24, 37, 50, 65, 80, and 98 □ were tested both in the presence and absence of MgCl_2_. Due to Mg^2+^-catalyzed RNA hydrolysis at high temperatures, RNA reactivity could not be measured at 98 □. Several reverse transcriptases, including SuperScript-II, Superscript-II with MnCl_2_, SuperScript-III, SuperScript-IV, TGIRT-III, MarathonRT, AMV, and MMLV, were tested in this study. Reaction conditions for each are listed in Table S1.

### Sequencing-based chemical mapping readout (MAP-seq)

To map terminations and mutations incorporated by different reverse transcriptases across from chemically modified TOD-S1 and TOD-S7 RNAs, we used the protocol described in ref^22^. Briefly, 5.0 pmol of RNA in 4 μL RNase free H_2_O was denatured by incubating at 95 °C for 2 min, and then cooling on ice for 1 min. Then 5.0 μL of 2X buffer (100 mM Na-HEPES, pH 8.0, and 20 mM MgCl_2_) was added, and the RNA was incubated at 50 °C for 30 min to fold. The RNA was modified by adding 1.0 μL of modifier (10.6 mg/mL of 1M7 in DMSO, 30 mg/ml of NMIA, or 1.25 % DMS in 8.75% EtOH). For no modification controls, 1.0 μL of RNase free H_2_O was added instead. Reactions were incubated at 24 °C for 15 min, and then quenched with 2 μL of 0.5 M Na-MES, pH 6.0 or β-mercaptoethanol for SHAPE or DMS modification, respectively. Modified RNA was then purified by ethanol precipitation. Reverse transcription and ligation of the second Illumina sequencing adapter was carried out as previously described.^22^ The RTB barcode assigned to each sample and the sequence of each RTB barcoded reverse transcription primer are listed in Supplemental File sequences.xlsx. For the second adapter ligation, P_TruSeqAdapt01_p and P_TruSeqAdapt02_p ligation primers were used for TOD-S1 and TOD-S7 constructs, respectively (see Supplemental File sequences.xlsx). Sequencing libraries were quantified by qPCR and sequenced on an Illumina MiSeq using a 150-cycle v3 chemistry kit per manufacturer’s instructions, with 35 and 121 cycles used for read 1 and read 2, respectively. Data were analyzed for termination events with MAPseeker software^22^ and for mismatch/deletion events with M2seq.py software (https://github.com/ribokit/M2seq)^42^.

### Chemical synthesis of modified RNAs

All chemically synthesized 41-mer TOD-S1 RNA molecules were generated at 40 nmol scale using the ABI-394 DNA/RNA synthesizer (Thermo Fisher Scientific) at the Stanford Protein and Nucleic Acid Facility (PAN). Through the process, synthesis and purification conditions avoided basic conditions that catalyze Dimroth rearrangement of m^1^A to m^6^A.^59^ For wild-type RNA molecules, UltraFast chemistry was used with Bz-A-CE, Ac-C-CE, Ac-G-CE, and U-CE phosphoramidites (Glen Research catalog numbers 10-3003-10, 10-3015-10, 10-3025-10, 10-3030-10, respectively). For RNAs containing m^1^A, UltraMild chemistry was used with Pac-A-CE, 1-Me-A-CE, Ac-C-CE, and iPr-Pac-G-CE phosphoramidites (Glen Research catalog numbers: 10-3000-10, 10-3501-95, 10-3015-10, 10-3021-10, 10-3030-10, respectively). (We could not chemically synthesize RNAs to test SHAPE-induced reverse transcriptase readthrough, as the exact 2’-hydroxyl adducts formed by SHAPE chemistry are not available as phosphoramidites.) Synthesized RNA was desalted with a Gel-Pak 1.0 desalting column (Glen Research) and concentrated with a SpeedVac (Thermo Fisher Scientific) on low heat. Concentrated RNA was checked for appropriate molecular weight with the Voyager-DE STR Biospectrometry Workstation (Applied Biosystems) and a MALDI-TOF mass spectrometer (Applied Biosystems). Quality-verified RNA was finally size selected using a denaturing 10% PAGE gel.

### ssRNA synthesis for NMR spectroscopy

12-mer RNA oligonucleotides were synthesized using a MerMade 6 Oligo Synthesizer employing 2′-tBDSilyl protected phosphoramidite (Bz-A-CE Phosphoramidite 10-3003-10, Ac-C-CE Phosphoramidite 10-3015-10, Ac-G-CE Phosphoramidite 10-3025-10, U-CE Phosphoramidite 10-3030-10 from ChemGenes) on 1 μmol standard synthesis columns (1000 Å) (BioAutomation). RNA oligonucleotides were synthesized with the option to leave the final 5□-protecting group (4,4□-dimethoxytrityl (DMT)) on. Synthesized oligonucleotides were cleaved from the 1 μmol column using 1 mL ammonia methylamine (1:1 ratio of 30% ammonium hydroxide and 30% methylamine) followed by 2-hour incubation at room temperature to allow base deprotection. The solution was then air-dried and dissolved in 115 μL DMSO, 60 μL TEA and 75 μL TEA-3HF, followed by 2.5 h incubation at 65 °C for 2′O-protecting group removal. Samples were then quenched with Glen-Pak RNA quenching buffer and loaded onto Glen-Pak RNA cartridges (Glen Research Corporation) for purification using the online protocol (http://www.glenresearch.com/). Samples were then ethanol precipitated, air-dried, dissolved in water and then buffer exchanged at least three times using a centrifugal concentrator (EMD Millipore) into the desired buffer.

### NMR experiments

All NMR experiments were collected on a 600 MHz or a 700 MHz Bruker NMR spectrometer equipped with an HCN cryogenic probe. Data were processed and analyzed using NMRpipe^60 1995^ and SPARKY^61 San Francisco^, respectively. Complete assignment experiments were obtained using 2D [^13^C, ^1^H] Heteronuclear Single Quantum Coherence (HSQC), 2D [^1^H, ^1^H] WATERGATE Nuclear Overhauser Effect Spectroscopy (NOESY, mixing time of 150 ms) experiments.

### DMS profiling an mutate-and-map-seq of HIV 3’UTR

For mutate-and-map-seq (and DMS-MaP-seq), the protocol in ref.^42^ was followed. RNA constructs for HIV 3’-UTR of reference genome HIV-1 NL4-3 were prepared without and without A_20_ inserted after the polyadenylation site (see Supplemental File sequences.xlsx). DNA templates (including prepended T7 promoter) were synthesized by PCR assembly based on primers designed by Primerize.^57^ PCR assembly was carried out using Phusion DNA polymerase as above, using annealing temperature of 64 °C for 35 cycles. DNA templates for mutate-and-map-seq were prepared with additional mutations by amplifying this DNA under error prone PCR conditions: 10 mM Tris-HCl, pH 8.3, 50 mM KCl, 4 mM MgCl_2_, 0.5 mM MnCl_2_, 1 mM dCTP, 1 mM dTTP, 0.2 mM dATP, 0.2 mM dGTP, 2 μM for forward and reverse primer, template 2 ng, Taq-DNA polymerase 5 units; PCR conditions were: 94 °C, 1 min-1 cycle; then 24 cycles of 94 °C for 30 sec, 64 °C for 1 min, 72 °C for 3 min; and 72 °C for 10 min for 1 cycle. DNA templates were purified by Qiagen PCR spin purification columns.

DNA templates were transcribed with T7 RNA polymerase (NEB) at 37 °C for 6 hrs, and RNA was purified by Agencourt Ampure XP beads supplemented with PEG-8000 (see above). In preparation for DMS chemical mapping, 12.5 pmol of RNA was heated at 95 °C for 2 min; cooled on ice for 1 min; then incubated in 300 mM Na-cacodylate, pH 7.0, 10 mM MgCl_2_ at 37 °C for 30 min at a final volume of 25 μL. Modification conditions used were 0.17 M of dimethyl sulfate at 37 °C for 6 min. RNAse free water was added instead as a no modification control. DMS reactions were quenched with 25 μL of β-mercaptoethanol. RNA was purified by ethanol precipitation in 10% v/v of 3 M sodium acetate. Purified RNA was resuspended in 7 μL of RNAse free water. 4.6 μL of the RNA sample was used to generated cDNA with SuperScript II reverse transcriptase with Mn^2+^ as the divalent ion to promote incorporation of mismatches or deletions across bypassed methylations. Reverse transcription reactions included 0.02 μM primer in final volume 12 μL; buffer of 25 mM Tris-HCl, pH 8.3, 75 mM KCl, 6 mM MnCl_2_, 5 mM DTT; and incubation of at 42 °C for 3 hrs. Reverse transcription reactions were stopped with 5 μL of 0.4 M sodium hydroxide at 90 °C for 3 min. Reactions were neutralized with acid quench (2 volumes 5 M NaCl, 2 volumes 2 M HCl, and 3 volumes of 3 M Na–acetate).

cDNA was purified with Agencourt Ampure beads supplemented with PEG-8000 (see above), and purified cDNA was resuspended in 12.5 μL of RNase free water. 2.5 μL of cDNA was used for PCR with Illumina adapters (see Supplemental File sequences.xlsx). We found it necessary to, perform emulsion PCR. For these reactions, we prepared an oil phase system composed of ABIL EM90, (Evonika) 80 μL, Triton X-100, 1.0 μL, and mineral oil, 1919 μL. Reactions combined 300 μL of this oil phase with 50 μL PCR mixture, composed of Phusion DNA polymerase, 1x Phusion buffer, 10 mM dNTPs, and 2 μM of each forward and reverse primer. We performed PCR at 98 °C, 30 sec-1 cycle; then 22 cycles of 98 °C for 10 sec, 64 °C for 30 sec, 72 °C for 30 sec; and 72 °C for 10 min for 1 cycle. The resulting dsDNA was purified by gel extraction and purification in Qiagen Qiaquick purification spin columns. The dsDNA was quantified with a Qubit instrument, with the HS dsDNA kit. A Miseq 600-cycle kit was used for sequencing; and data were analyzed with the M2seq.py pipeline described in ref^42^.

## Data accessibility

Data for reverse transcribed cDNAs are deposited at the RNA Mapping Database^11^ (https://rmdb.stanford.edu) with the following accession IDs:

|  |  |
| --- | --- |
| TOD-S1 RNA, 1M7 (SHAPE) across different solution conditions | TODS1_1M7_0001 |
| TOD-S1 RNA, DMS across different solution conditions | TODS1_DMS_0001 |
| TOD-S7 RNA, 1M7 (SHAPE) across different solution conditions | TODS7_1M7_0001 |
| TOD-S7 RNA, DMS across different solution conditions | TODS7_DMS_0001 |
| TOD-S1 RNA, 1M7 (SHAPE), vary RTases | TODS1_1M7_0002 |
| TOD-S1 RNA, DMS, vary RTases | TODS1_DMS_0002 |
| TOD-S7 RNA, 1M7 (SHAPE), vary RTases | TODS7_1M7_0002 |
| TOD-S7 RNA, DMS, vary RTases | TODS7_DMS_0002 |
| TOD-S7 RNA; 1M7, NMIA, DMS; vary RTases; terminations read out by Illumina sequencing | TODS7_TRM_0001 |
| TOD-S7 RNA; 1M7, NMIA, DMS; vary RTases; mismatches/deletions read out by Illumina sequencing | TODS7_MUT_0001 |
| Syn41 RNA, chemically synthesized with m<sup>1</sup>A at different positions, no further modification, varying reverse transcriptases | SYN41_M1A_0001 |
| HIV 3' UTR, no A<sub>20</sub> tail, DMS | HIV3PR_DMS_0001 |
| HIV 3' UTR, no A<sub>20</sub> tail, no modifier control | HIV3PR_NMD_0001 |
| HIV 3' UTR, no A<sub>20</sub> tail, DMS, transcribed off error-prone PCR DNA (mutate-and-map-seq) | HIV3PR_DMS_0002 |
| HIV 3' UTR, no A<sub>20</sub> tail, no modifier control, transcribed off error-prone PCR DNA | HIV3PR_NMD_0002 |
| HIV 3' UTR, with A<sub>20</sub> tail, DMS | HIV3PR_DMS_0003 |
| HIV 3' UTR, with A<sub>20</sub> tail no modifier control | HIV3PR_NMD_0004 |
| HIV 3' UTR, with A<sub>20</sub> tail, DMS, transcribed off error-prone PCR DNA (mutate-and-map-seq) | HIV3PR_DMS_0004 |
| HIV 3' UTR, with A<sub>20</sub> tail, no modifier control, transcribed off error-prone PCR DNA | HIV3PR_NMD_0004 |

