## Supplemental Tables and Figures for "Anomalous reverse transcription through chemical modifications in polyadenosine stretches"

### Supplemental Information for: Anomalous reverse transcription through chemical modifications in polyadenosine stretches

Wipapat Kladwang[1], Ved T. Topkar[2], Bei Liu[3], Tracy L. Hodges[4], Sarah C. Keane[4,5], Hashim al-Hashimi[3,6], Rhiju Das[1,2,7,*]

[1]Department of Biochemistry, Stanford University School of Medicine, Stanford CA 94305
[2]Biophysics Program, Stanford University, Stanford CA 94305
[3]Department of Biochemistry, Duke University School of Medicine, Durham NC 27710
[4]Biophysics Program, University of Michigan, Ann Arbor MI 48109
[5]Department of Chemistry, University of Michigan, Ann Arbor MI 48109
[6]Department of Chemistry, Duke University School of Medicine, Durham NC 27710
[7]Department of Physics, Stanford University, Stanford CA 94305

**Table S1. Reverse transcriptase conditions tested for poly(A) bypass**

| No. | Reverse-transcriptase | Buffer* | Reaction conditions | Enzyme source | Part number | Stock | Units/reaction |
|---|---|---|---|---|---|---|---|
| 1 | SuperScript-II | 25 mM Tris-HCl- pH 8.3, 75 mM KCl, 3 mM $MgCl_2$, 5 mM DTT** | 42 °C | Thermo Fisher Scientific | 18064071 | 200 U/$\mu$L | 20 Units |
| 2 | SuperScript-II + $Mn^{2+}$ | 25 mM Tris-HCl- pH 8.3, 75 mM KCl, 6 mM $MnCl_2$, 5 mM DTT | 42 °C | Thermo Fisher Scientific | 18064071 | 200 U/$\mu$L | 20 Units |
| 3 | SuperScript-III | 25 mM Tris-HCl- pH 8.3, 75 mM KCl, 3 mM $MgCl_2$, 5 mM DTT** | 48 °C | Thermo Fisher Scientific | 18080085 | 200 U/$\mu$L | 20 Units |
| 4 | SuperScript-IV | 50 mM Tris-HCl- pH 8.3, 4 mM $MgCl_2$, 50 mM KCl, 10 mM DTT** | 50 °C | Thermo Fisher Scientific | 18090050 | 200 U/$\mu$L | 20 Units |
| 5 | TGIRT-III | 50 mM Tris-HCl-pH 8, 75 mM KCl, 3 mM $MgCl_2$ 5 mM DTT | 57 °C | InGex, LLC | TGIRT50 | 280 U/$\mu$L | 140 Units |
| 6 | Marathon RT | 50 mM Tris-HCl- pH 8.5, 100 mM KCl, 2 mM $MgCl_2$ 10 mM DTT | 42 °C | Courtesy of Dr. A. Pyle, Yale University | n/a | 5 $\mu$M | 4.6 pmol |
| 7 | AMV | 25 mM Tris-HCl- pH 8.3, 100 mM KCl, 5 mM $MgCl_2$, 2 mM DTT** | 42 °C | AMS Biotechnology Ltd. | AMS.AMV007-1 | 20 U/$\mu$L | 20 Units |
| 8 | MMLV | 50 mM Tris-HCl- pH 8.3, 75 mM KCl, 3 mM $MgCl_2$, 5 mM DTT** | 42 °C | Thermo Fisher Scientific | AM2043 | 100 U/$\mu$L | 20 Units |

*60 min reactions in 5 $\mu$L volumes

**Manufacture-supplied buffer was used.

**Table S2. RTB barcoded reverse transcription primers for MAP-seq cDNA library preparation**

| **RNA/ modification** | **SS-II** | **SS-II + Mn** | **SS-III** | **SS-IV** | **TGIRT** | **Marathon** | **AMV** | **MMLV** |
|---|---|---|---|---|---|---|---|---|
| TOD-S1-No-Modification | RTB000 | RTB001 | RTB002 | RTB003 | RTB004 | RTB005 | RTB006 | RTB007 |
| TOD-S1-1M7 | RTB008 | RTB009 | RTB010 | RTB011 | RTB012 | RTB013 | RTB014 | RTB015 |
| TOD-S1-NMIA | RTB016 | RTB017 | RTB018 | RTB019 | RTB020 | RTB021 | RTB022 | RTB023 |
| TOD-S1-DMS | RTB024 | RTB025 | RTB026 | RTB027 | RTB028 | RTB029 | RTB030 | RTB031 |
| TOD-S7-No-Modification | RTB000 | RTB001 | RTB002 | RTB003 | RTB004 | RTB005 | RTB006 | RTB007 |
| TOD-S7-1M7 | RTB008 | RTB009 | RTB010 | RTB011 | RTB012 | RTB013 | RTB014 | RTB015 |
| TOD-S7-NMIA | RTB016 | RTB017 | RTB018 | RTB019 | RTB020 | RTB021 | RTB022 | RTB023 |
| TOD-S7-DMS | RTB024 | RTB025 | RTB026 | RTB027 | RTB028 | RTB029 | RTB030 | RTB031 |

**Table S3. RTB barcoded reverse transcription primers for mutate-and-map-seq/DMS-MaP-seq experiments on HIV 3´-UTR RNAs.**

| Number | RNA* | RTB primer | |
|---|---|---|---|
| | | No mod | High DMS |
| 1 | P4P6.noHP-Phu | RTB000 | RTB001 |
| 2 | P4P6.noHP-Taq | RTB002 | RTB003 |
| 3 | NL4-3 with polyA tail -Phu | RTB004 | RTB005 |
| 4 | NL4-3 with polyA tail -Taq | RTB006 | RTB007 |
| 5 | WO-A-Phu | RTB008 | RTB009 |
| 6 | WO-A.noHP-Taq | RTB010 | RTB011 |

* Phu – standard PCR preparation with Phusion polyerase, Taq – error-prone PCR with Taq polymerase and $Mn^{2+}$, to introduce mutations for mutate-and-map-seq analysis, NL4-3 – HIV 3´ UTR sequence from NL4-3 genome, WO-A – no $A_{20}$.

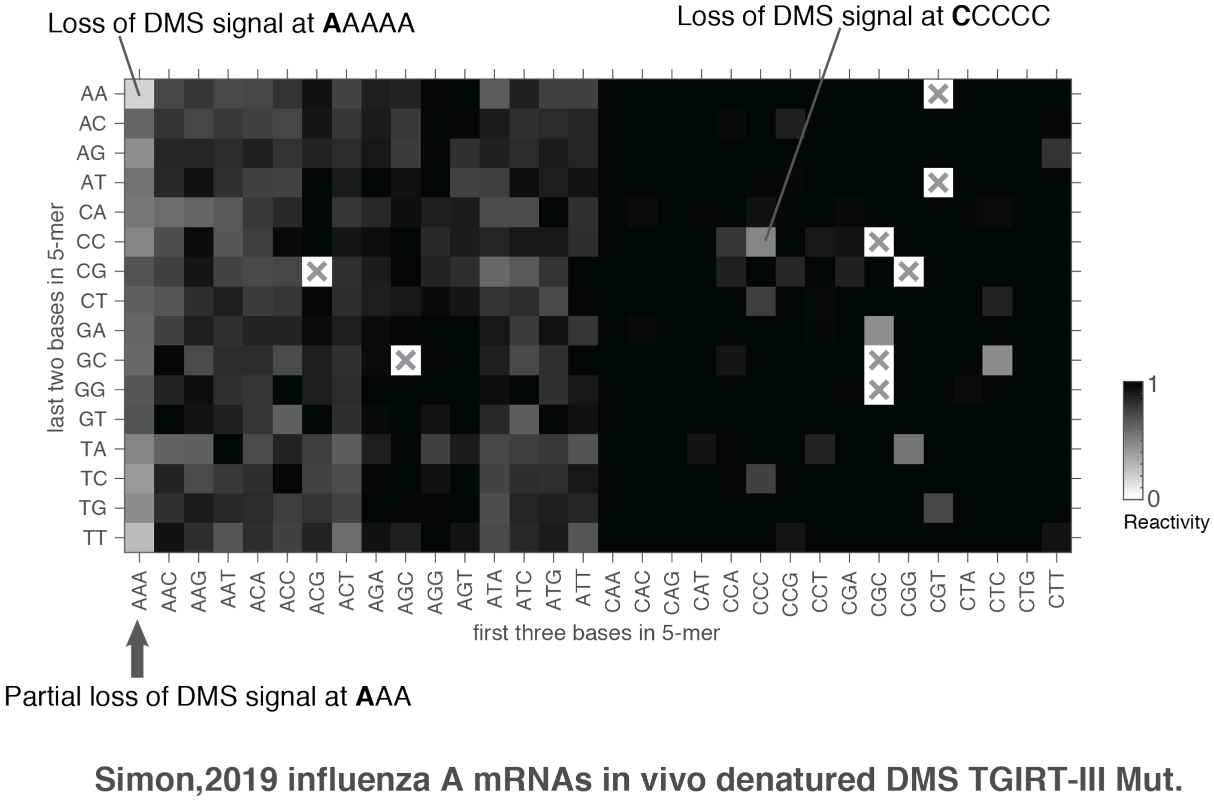


**Figure S1. Apparent DMS signals for nearly all 5-mer sequences read out through mutational profiling.** Data are for denatured influenza A mRNAs in (Simon,2019), averaged for each occurrence of the 5-mer. Reactivities are normalized to mean DMS reactivity at all 5-mers for which data are available. X's mark 5-mers for which data were not available. Loss of DMS signals at AAA triplets, and stronger loss at AAAAA and CCCCC are marked.

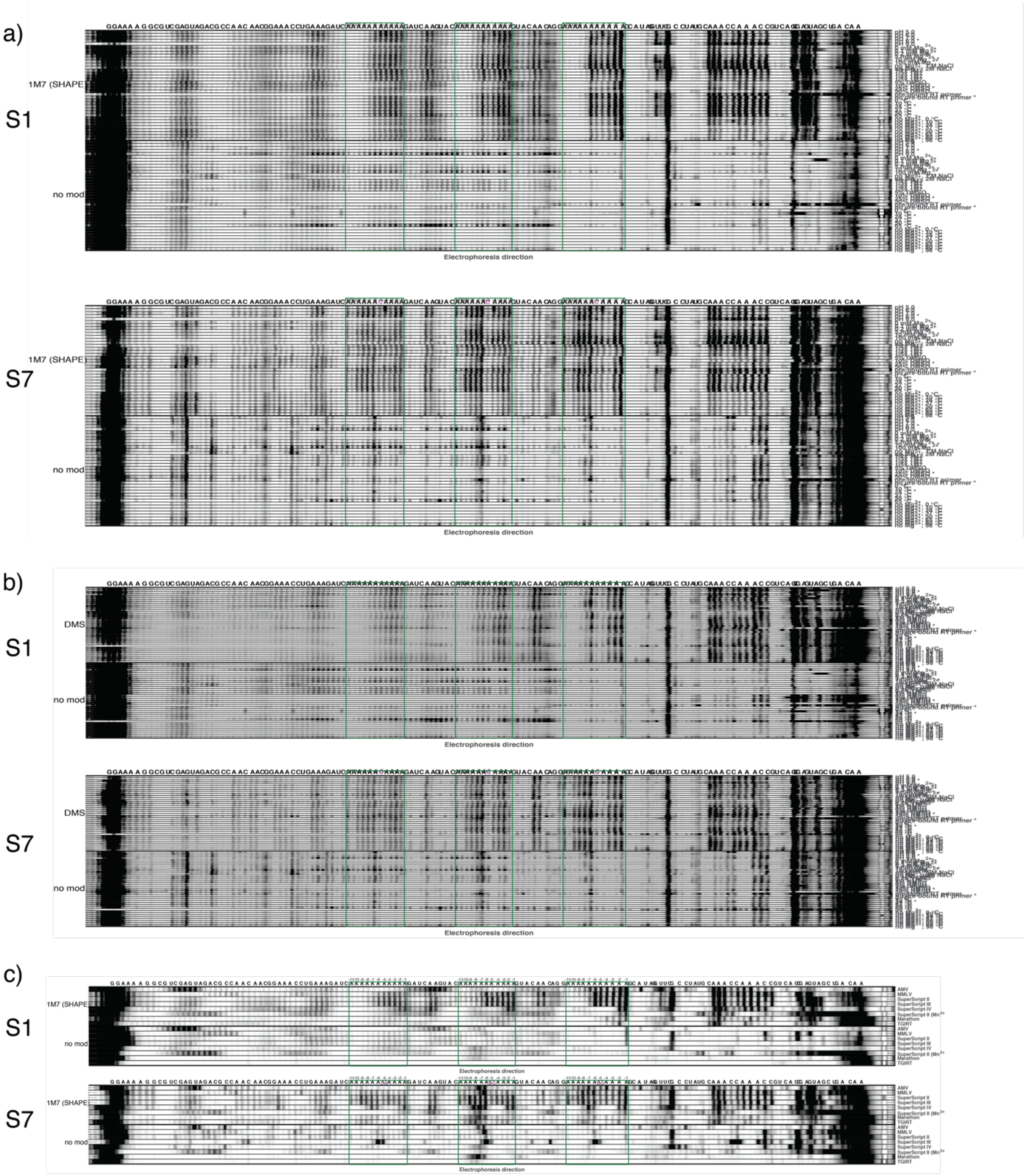


**Figure S2. Capillary electropherograms for TOD-S1 and TOD-S7 experiments.** (a,b) Variation of solution conditions during (a) SHAPE (1M7) or (b) DMS mapping, with no modification control data acquired side-by-side shown underneath. (c) Variation of reverse transcriptases after SHAPE (1M7) mapping. In each panel, stretches of $A_{11}$ of AAAAAACAAAA are outlined in green boxes.

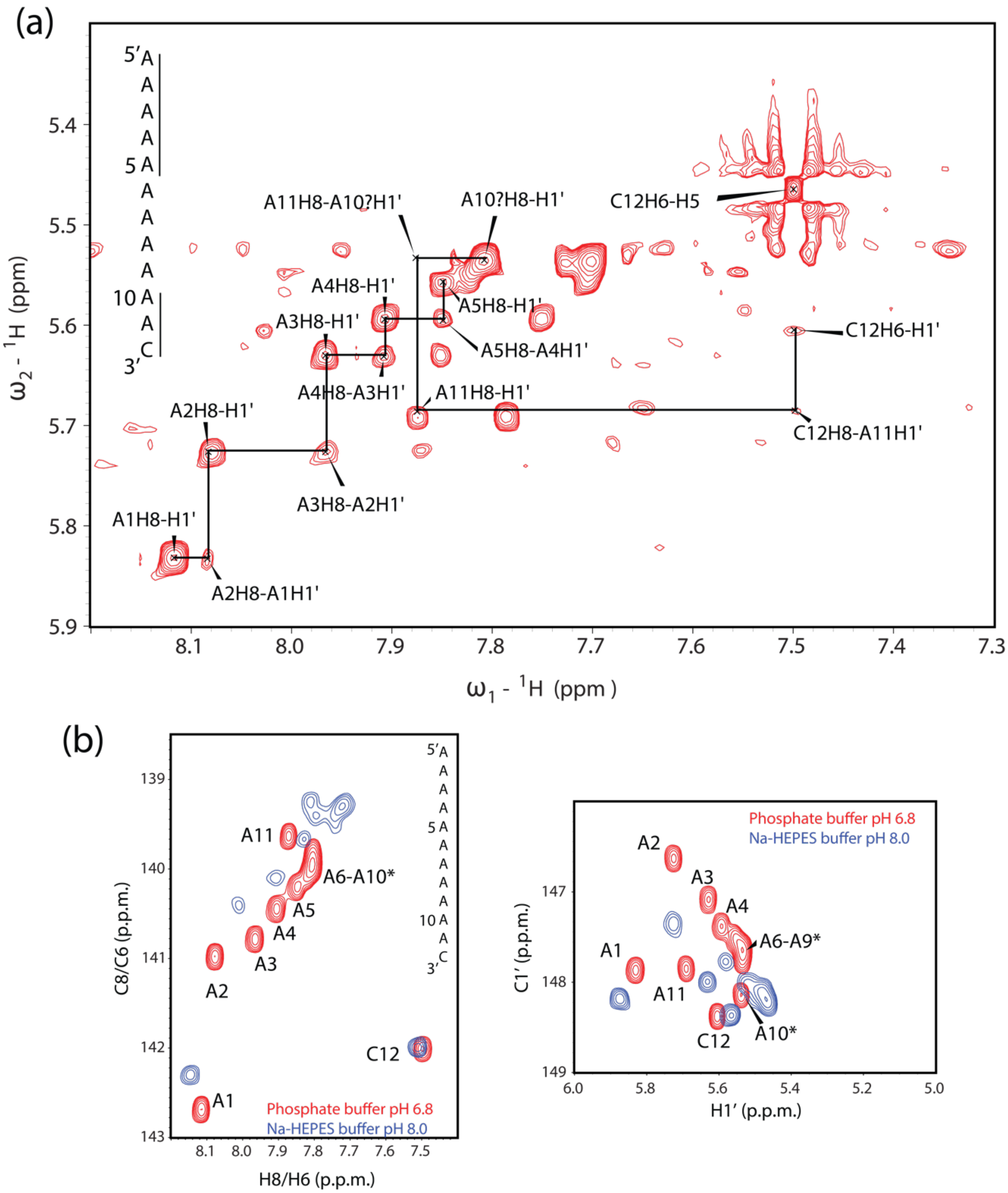


**Figure S3. NMR spectra of $A_{11}C$ RNA test for unusual structures.** (a) 2D NOESY spectra of $A_{11}C$ RNA (5'-AAAAAAAAAAAC-3') in phosphate buffer (25 mM NaCl, 15 mM sodium phosphate, pH 6.8, 25°C; RNA concentration of 1 mM) enable chemical shift assignment. (b) 2D [$^{13}C$,$^{1}H$] HSQC spectra of C8 and H8 atoms within aromatic bases and C1´ and H1´ within riboses for A11C RNA in Na-HEPES buffer that matches our chemical mapping experiments (blue, 50 mM Na-HEPES, pH 8.0, 10 mM $MgCl_2$, T = 25°C; RNA concentration, 1 mM) and in phosphate buffer more standard for NMR (red). Chemical shifts and NOE cross-peaks remain consistent with a dominant A-form like structure for the RNA across buffer conditions.

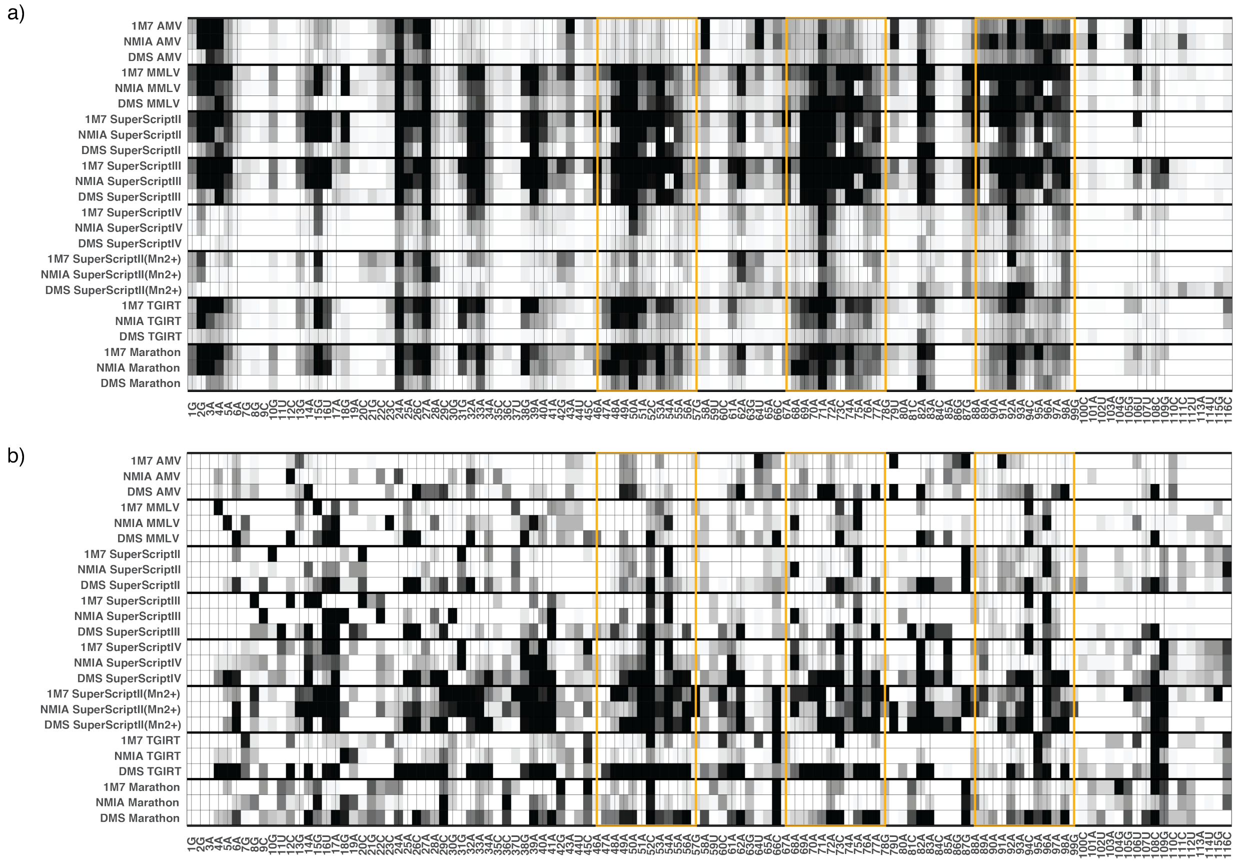


**Figure S4. Illumina sequencing confirms bypass at poly(A) stretches depending on reverse transcriptase**. Fractions of products (a) terminated or (b) inducing mismatches/deletions in TOD-S7 RNA, after background subtraction. Thin black vertical lines mark bases designed to be unpaired in the molecules. Gold rectangles highlight repeats of AAAAAACAAAA stretch.

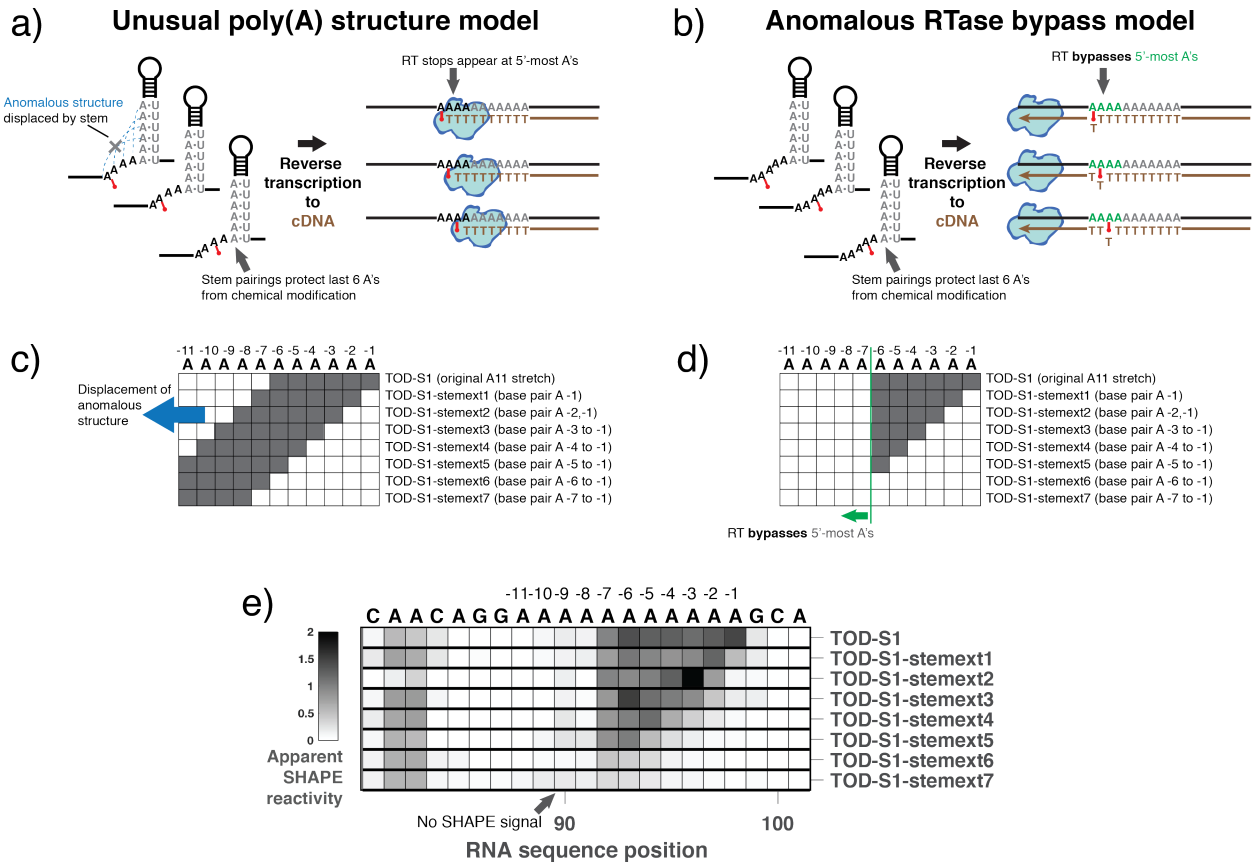


**Figure S5. Templates with additional designed base pairs further implicate the RTase bypass model.** (a-b) TOD-S1-ext RNAs introduce stretches of U's near each of the three 11-adenosine stretches and were designed to extend the RNA's three helical stems to include additional A-U pairs. (a,c) in the poly(A) structure model, if A's were taken out of an anomalous structure and into these canonical base pairs, the poly(A) anomalous structure (a) and associated chemical modification signals (c) should shift 5'. (b,d) In the 'reverse transcriptase bypass' model, the chemical modification signals at newly paired A's should disappear, but no new cDNA products corresponding to new chemical modifications at other A's should appear. (e) The experimental data match the predictions of the RTase bypass model (d) and disfavor the poly(A) structure model (c).

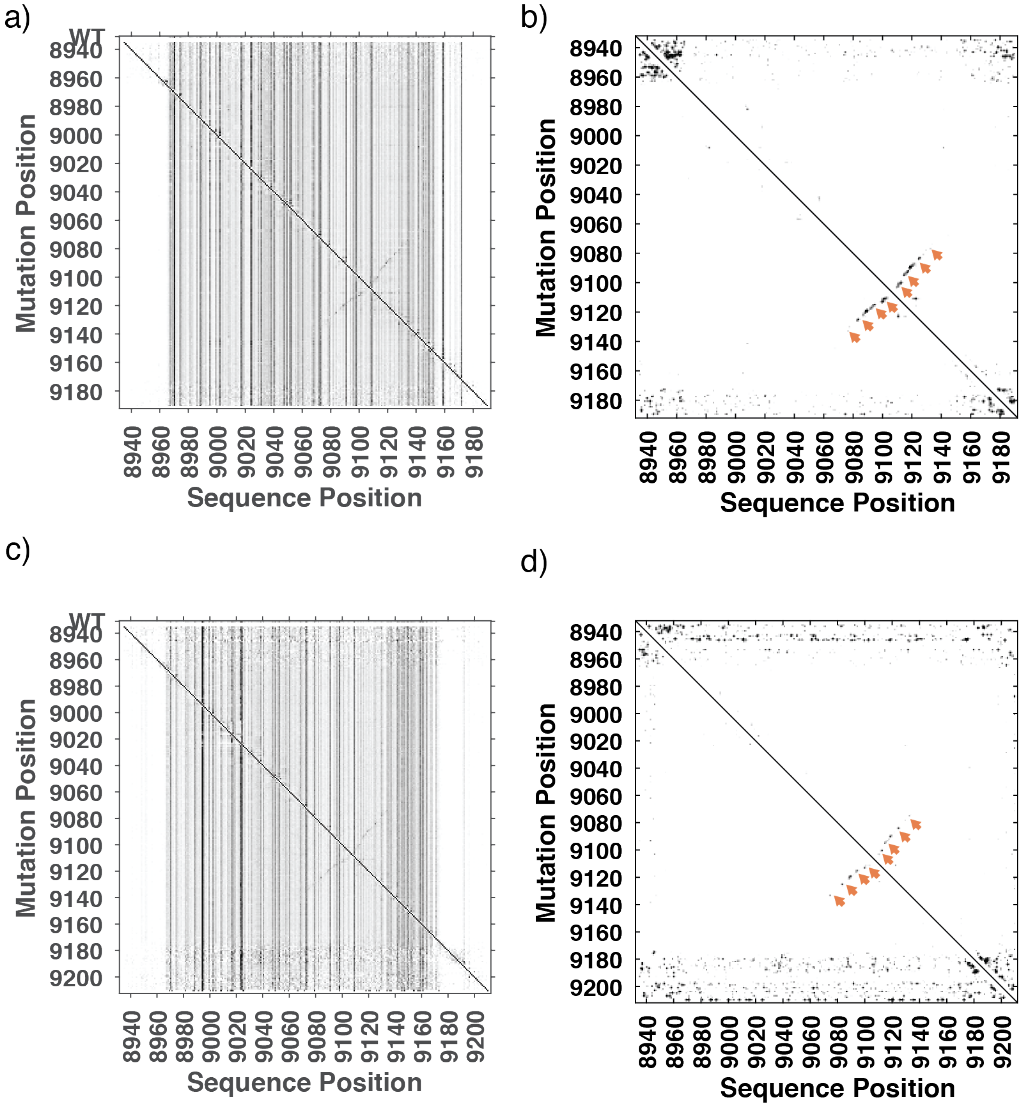


**Figure S6. Mutate-and-map-seq experiments on HIV-1 3'-UTR.** Numbering is for HIV-1 NL4-4 reference genome. Constructs stop at polyadenylation site (a,b) or are further extended by $(A)_{20}$ (c,d); each are then extended by a 20-nt primer binding site. (a,c) Two-dimensional M2-seq signal shows mutation rate (primarily reflecting DMS reactivity) at each sequence position given detection of one mutation (primarily deletion/mismatches introduced by error-prone PCR) at other sites. (b,d) Z-score maps (smoothed) based on M2-seq signals; arrows mark stems that are automatically detected with the M2-net algorithm and are also clear visually. All four stems comprising the HIV-1 3' TAR hairpin are detected, in both data sets. See Cheng, C.Y., Kladwang, W., Yesselman, J.D., and Das, R. (2017), "RNA structure inference through chemical mapping after accidental or intentional mutations", *Proceedings of the National Academy of Sciences U.S.A.* 114 (37) : 9876 – 9881.
